# A kinetic model of antigen-dependent IgG oligomerization and complement binding

**DOI:** 10.1101/2025.02.04.635266

**Authors:** Jürgen Strasser, Nikolaus Frischauf, Lukas Schustereder, Andreas Karner, Sieto Bosgra, Aran F. Labrijn, Frank J. Beurskens, Johannes Preiner

**Affiliations:** University of Applied Sciences Upper Austria, Linz, Austria; Genmab, Utrecht, Netherlands

**Keywords:** IgG hexamers, IgG oligomerization, classical complement pathway, IgG subclasses, C1, kinetic model, complement mediated lysis

## Abstract

The classical complement pathway (CCP), an important branch of the mammalian immune system, is initiated through multivalent binding of complement protein C1 to Immunoglobulin G (IgG) antibody oligomers assembled on the surface of pathogens, infected or malignant cells, culminating in the formation of the membrane attack complex (MAC) and subsequent cell lysis. IgG oligomers can further engage immune effector cells through Fcγ receptors or complement receptors, facilitating antibody-dependent cellular cytotoxicity (ADCC) and phagocytosis (ADCP). Detailed knowledge of the factors that drive IgG oligomerization is thus vitally important to establish and improve IgG based therapies. We here focus on the kinetics of antigen-dependent IgG oligomerization and develop a comprehensive model capable of predicting oligomer formation as a function of IgG concentration, antigen density, IgG subclass and Fc point mutants, as well as the presence of Fc-binding and thus oligomerization-inhibiting factors such as staphylococcal protein A (SpA). We characterize the underlying molecular interactions in single molecule force spectroscopy (SMFS) and grating coupled interferometry (GCI) experiments. By fitting experimental data from high-speed atomic force microscopy (HS-AFM) experiments, we further quantify key rate constants and thermodynamic parameters, including free energy changes associated with oligomerization and apply the model to predict complement-mediated lysis in liposomal vesicle-based assays. The presented mechanistic framework may serve as a basis for optimizing antibody engineering and pharmacokinetic/pharmacodynamic modeling in the context of immunotherapies exploiting the CCP.

## INTRODUCTION

Multivalent interactions are ubiquitous in biological systems as they enable complex molecular recognition and signaling processes. These interactions occur when multiple binding sites on a molecule or cell surface engage with multiple counterparts, leading to enhanced specificity and strength of the overall interaction. A prime example of multivalent interactions that trigger complex immunological effector functions is the initiation of the classical complement pathway (CCP) through the antigen-dependent oligomerization of Immunoglobulin G (IgG) antibodies and the subsequent recognition and activation of the multivalent zymogen C1 by IgG tetramers to hexamers^1–5^. Multiple IgG Fab – antigen, IgG Fc – Fc, and IgG – C1 interactions thereby act together to form a complex that launches an amplifiable cascade of soluble zymogens abundant in blood and other extracellular fluids that may lead to insertion of a lytic pore (membrane attack complex; MAC) into the target cell membrane^6^ (complement dependent cytotoxicity; CDC). Alternatively, membrane-bound IgG oligomers or downstream CCP components activate Fcγ receptor-or complement receptor-expressing effector cells such as natural killer (NK) cells, macrophages or neutrophils that ultimately kill the IgG-opsonized target cell (antibody dependent cellular cytotoxicity/phagocytosis; ADCC/ADCP)^7,8^.

The four human IgG subclasses significantly differ in their potency to activate the CCP under otherwise identical conditions, which we could recently attribute to their individual ability to form IgG tetramers and higher order oligomers that are needed to activate C1^5^. However, IgG oligomerization can be enhanced in all IgG subclasses by point mutations at the Fc domain, such as E430G, leading to an improved abundance of higher oligomers and thus complement activation, especially at lower antigen densities where wild type IgGs are less efficient in forming higher oligomers or fail entirely. We have previously developed a kinetic antigen-dependent oligomerization model for functionally monovalent IgG1 and applied it to respective binding curves recorded in real-time quartz crystal microbalance (QCM) experiments^4^. However, a comprehensive model valid for functionally bivalent (natural) IgGs capable of predicting oligomerization in dependence of IgG concentration, antigen density, IgG subclass/ mutant variant and Fc-binding factors such as staphylococcal protein A (SpA)^9^ has not been established yet. In this study, we biophysically characterize the molecular interactions, kinetic rate constants and thermodynamic properties underlying the initial steps of the IgG-mediated CCP. We present a mechanistic framework for predicting IgG oligomerization, subsequent C1 binding, and complement mediated lysis in a liposomal vesicle-based assay, considering the impact of Fc point mutants, IgG subclasses, or Fc – Fc interaction inhibiting factors. The presented kinetic model may thus serve as a basis for optimizing protein-engineering of antibody formats and as a starting point for comprehensive pharmacokinetic/pharmacodynamic (PK/PD) modelling in establishing dosing schemes for therapeutic IgGs in immunotherapies.

## RESULTS AND DISCUSSION

### A mechanistic model of IgG oligomerization

We here extend our previous model^4^ to functionally bivalent IgGs (Figure 1) and apply it to IgG oligomer distributions obtained from HS-AFM experiments. The model for monovalent IgGs consists of two different oligomerization pathways that differ in the sequence of binding events. In the ‘lateral pathway’, IgGs first bind monovalently to surface antigens and subsequently form higher oligomers through Fc – Fc interactions enabled by lateral diffusion driven encounters. In the ‘vertical pathway’, on the other hand, antigen-bound IgGs recruit additional IgGs from solution via Fc – Fc interactions. Lateral oligomerization was experimentally observed only for monovalently epitope-bound IgGs but not for bivalently epitope-bound IgGs, since steric constraints did not allow for Fc – Fc interactions between neighboring IgGs^3^. Owing to the antigen densities employed in our model system, functionally bivalent IgGs quickly re-establish bivalent binding after dissociation of a single Fab domain so that contributions from the lateral pathway to the overall oligomerization can be neglected. For the same reason, the lower branch of the vertical pathway that includes only monovalently epitope-bound IgGs (Figure 1B) does not significantly contribute to the overall oligomerization under these conditions and is thus only considered here for the sake of completeness as it likely plays a more important role at antigen densities where bivalent binding is less likely.

**Figure 1.**
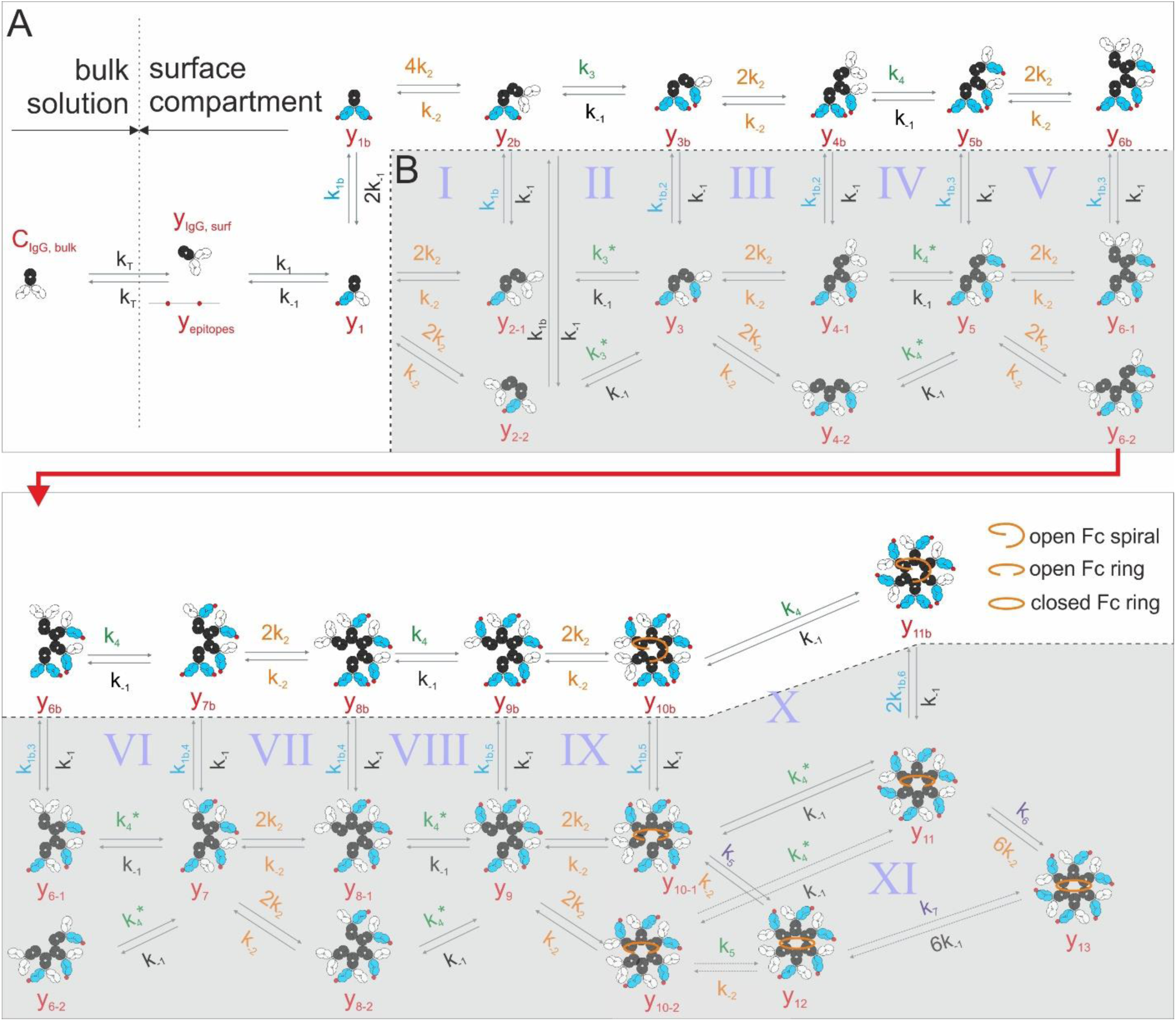
Kinetic model of antigen-dependent IgG oligomerization. **(A)** Upper branch of the vertical oligomerization pathway. **(B)** Lower branch of the vertical pathway. Fc domains, antigen-unbound and antigen-bound Fab domains are colored black, white and blue, respectively.

We assume that the kinetic rates governing the Fc – Fc interactions (k_2_ / k_-2_) are independent of the binding valency to surface epitopes (i.e. that no conformational change occurs upon binding), however, the possible directions of growth of an oligomer might differ for these two situations. While unoccupied Fc binding sites within IgG oligomers with strictly monovalently epitope bound IgGs have likely largely identical distances/orientations with respect to antigenic surface (e.g. y_3_, y_5_ etc.) and can thus likely grow bi-directionally (e.g. via y_4-1_ or y_4-2_, y_6-1_ or y_6-2_ etc.), the two free Fc binding sites on IgG oligomers that include a bivalently bound IgG (e.g. y_3b_, y_5b_ etc.) likely exhibit a less symmetric configuration (compare also open Fc spiral/ring sketched in Figure 1) resulting in a preferred direction of growth that is defined upon binding of the first IgG to the nucleating bivalently bound IgG (e.g. via y_4b_, y_6b_ etc.). Consequently, we assume different rates for these steps, where a transiently Fc-bound IgG may establish an interaction with a surface epitope (k_3_*/ k_3_), a process that critically depends on the motional degree of freedom of the respective complex relative to surface epitopes. Due to the principle of microscopic reversibility, the upper and the lower branches need to be connected all the way through the formation of IgG hexamers and the respective rates must meet constraints (I – X in Figure 1) of the type

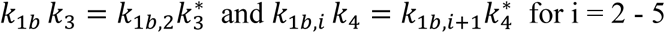

which can be summarized to

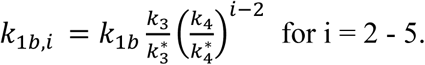

Furthermore, condition XI must be met, resulting in

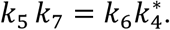

These constrains guarantee that the Gibb’s free energies accumulated through different pathways connecting two arbitrary states are equal. Accounting for mass transport limitations in our experimental system, we implemented a two-compartment model to mimic the IgG concentration differences in the bulk solution and close to the antigenic surfaces where IgG binding results in a partial depletion of solution phase IgGs, thus lowering the effective concentration in the surface compartment^4,10^. As mentioned above, we only consider the upper branch (white area in Figure 1) of the vertical pathway since at the antigen densities used in our model system the lower branch states are not significantly populated. Additionally, since we cannot distinguish hexameric states y_11-13_ from state y_11,b_ in our HS-AFM experiments, the rates k_5-7_ cannot be determined from the fits. The resulting model thus yields the following set of rate equations:

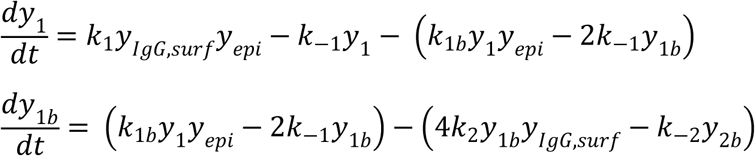

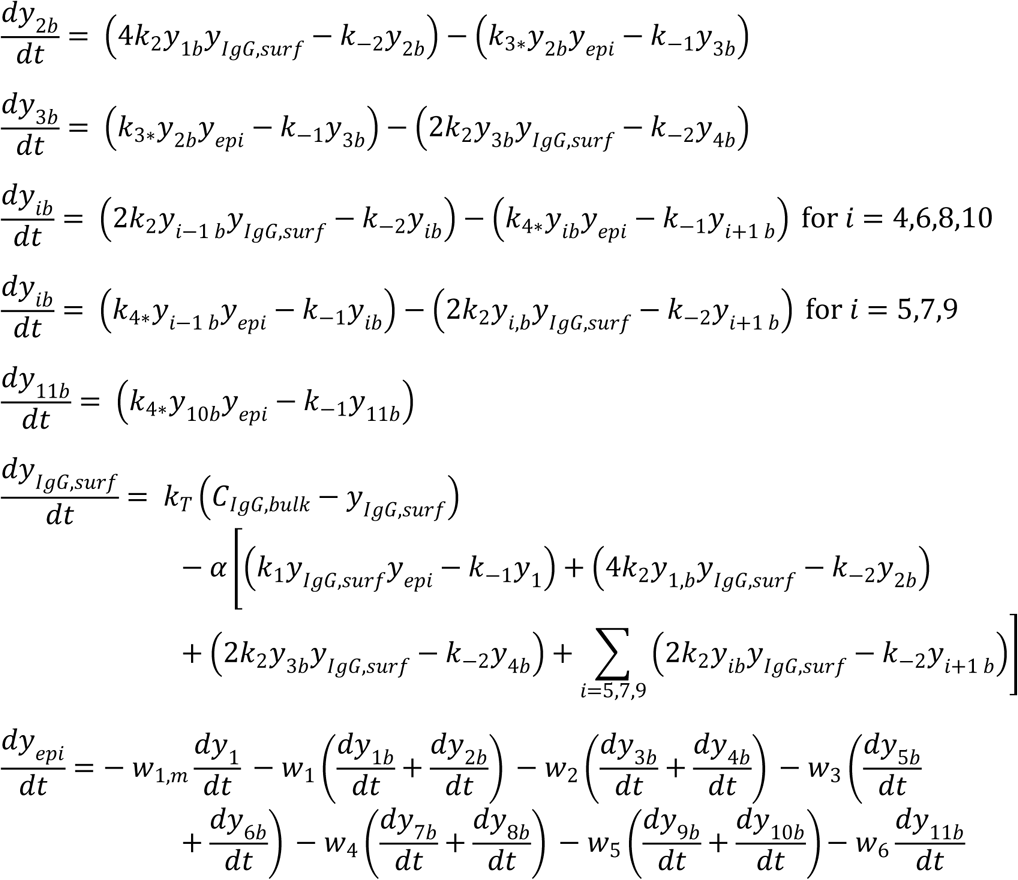

α = 1⁄(6.022 · 10^8^) μ is the factor by which the concentration in the surface compartment changes through the association/dissociation of 1 IgG/µm^2^ assuming a surface compartment size of 1 µm (perpendicular to the surface; 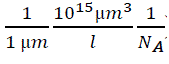) and being instantaneously well-mixed. The parameters *w_i_* account for epitopes (surface density [epi]) that are unbound but also unavailable as they are screened by (hidden below) other IgGs bound to neighboring epitopes and are given by^4^:

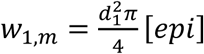

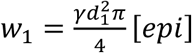

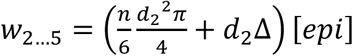

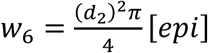

A single monovalent IgG screens a circular patch of diameter d_1_=8.8 nm ^11^, a bivalently bound IgG screens a circular patch of diameter 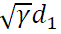 with γ = 1.6 ^11^, hexamers screen a circular patch of diameter d_2_=23 nm ^2^, and dimers to pentamers screen the respective fragment of a hexamer plus an additional area (*d*_2_Δ, with Δ=2.9 nm) accounting for overlap effects at the rim (along the Fc portions) of these oligomers. We have further extended the model to include the interaction of IgGs with a competitor that binds to IgG Fc domains such as the B domain of SpA (SpA-B_AA_)^9^ and thereby blocks Fc-Fc interactions (Figure S1; SI Text), an experimental strategy we apply in SMFS and HS-AFM experiments as shown below.

### Determination of kinetic parameters governing the Fc – Fc interactions

To reduce the number of floating parameters that need to be adjusted when fitting the model to HS-AFM data, we independently determined the association/dissociation rate constants governing the Fc – Fc interaction between antigenic-epitope bound and unbound IgGs in atomic force microscopy (AFM)-based single molecule force spectroscopy (SMFS) experiments^12^ (Figure 2A). By measuring dissociation forces between individual IgGs, these experiments allow for the characterization of transient/weak interactions which cannot be readily detected using conventional ensemble methods, as proven previously when we used this technique to characterize Fc – Fc interactions of the IgG1 subclass in solution in the absence of an antigenic surface^4^. We prepared supported lipid bilayers (SLB) containing 5% dinitrophenyl (DNP)-labeled lipids^3,4^ on glass coverslips and incubated them with a defined IgG concentration of a certain subclass as well as the E430G point mutant of IgG1-DNP, IgG1-DNP-E430G, to generate sparsely distributed IgG oligomers. These IgG oligomer-decorated SLBs were than examined in SMFS experiment using a cantilever tip to which a corresponding DNP-unspecific IgG or mutant variant was covalently attached through a flexible polyethylene glycol (PEG) linker, providing the tip-linked IgG with a high motional freedom mimicking the situation in solution^13^. For each IgG variant, force vs. distance (FD) curves were recorded at a wide range of loading rates revealing characteristic nonlinear force spectra in the pN regime (Figure 2B). For IgG1, we performed three different versions of this experiment comparing homotypic IgG1 – IgG1, IgG1-E430G – IgG1-E430G and IgG1 – IgG1-Fc interactions. Force spectra of IgG1 – IgG1 and IgG1 – IgG1-Fc interactions largely overlapped, indicating that indeed only intermolecular contacts between the Fc domains (and not Fab domains) contribute to the binding. Analysis according to the Bell-Evans model^12^ revealed dissociation rate constants (k_off_) and a structural parameter (x_b_) describing the distance from the bound to the transition state in the Gibbs free energy landscape (Figure 2D). We found no significant difference in *k*_off_ for these three experiments (1.5 (0.7, 2.6) s^−1^ vs. 1.8 (0.9, 3.0) s^−1^ vs. 1.3 (0.4, 2.7) s^−1^; Figure 2C, upper panel) confirming that the E430G point mutation does not change the strength of the Fc – Fc interaction. We did observe a larger *xβ*, however, which may be interpreted as a difference in flexibility of IgG1-E430G during the pulling experiment (Figure 2C, middle panel). While *xβ* is likely to depend on the exact attachment point of the linker molecule and should therefore be interpreted with caution, this difference could also be rationalized considering that the E430G point mutation removes a salt bridge stabilizing the CH2−CH3 interface packing and is thus likely to increase the flexibility of the Fc domain^4,5,7^. IgG2 (1.1 (0.1, 4.5) s^−1^) and IgG4 (0.55 (0.02, 2.84) s^−1^) exhibited dissociation rate constants comparable to IgG1, but IgG3 had a significantly lower *k_off_* = 0.07 (0.01, 0.2) s^−1^. Recent structural data suggests that additionally to IgG3 Fc – Fc interactions, IgG3-Fab_2_s form homotypic contacts resulting in a regular, 2D array formation below the hexameric Fc platform on sufficiently fluid lipid membranes^14^. To check for potential contributions of such contacts to the interactions detected in our SMFS experiments, we performed control experiments on IgG3-DNP oligomer decorated DNP-SLBs but using IgG3-Fab_2_-CD52 and IgG3-Fc domains instead of full length IgG3 linked to the AFM tip. The resulting dissociation rate constants of 2.6 (0.8, 5.6) s^−1^ (IgG3-Fab_2_) and 0.04 (1 x 10^−10^, 1.5) s^−1^ (IgG3-Fc) suggest that the main contributions of the interactions between the full length IgG3s stems from homotypic Fc – Fc interactions (Figure S2). The specificity of these interactions was confirmed in two control experiments, involving either a bare DNP-SLB or the presence of 4.5 µM SpA-B_AA_ that was shown to effectively block IgG Fc – Fc interactions through binding to the Fc – Fc interfaces (except for IgG3)^9^ (Figure 2C, lower panel). In both types of experiment, binding probability is markedly reduced confirming the interactions to be specific Fc – Fc interactions. We further estimated the association rate constants of the respective interactions from the dependency of the binding probability (i.e., the probability to observe a dissociation event in a given set of FD curves) on the encounter time of tip– and surface-bound IgGs during which a complex can be formed^4,15^. Varying this time span and evaluating the corresponding binding probabilities yields typical saturation curves (Figure 2E), resulting in 1–4 x 10^5^ M^-1^s^−1^ for all IgG variants, albeit with relatively large confidence intervals so that the values can only be regarded as a vague order of magnitude (Figure 2F, upper panel), which is also true for the equilibrium dissociation constants calculated using K_D,Fc_ = k_off_/k_on_ (Figure 2F. lower panel).

**Figure 2.**
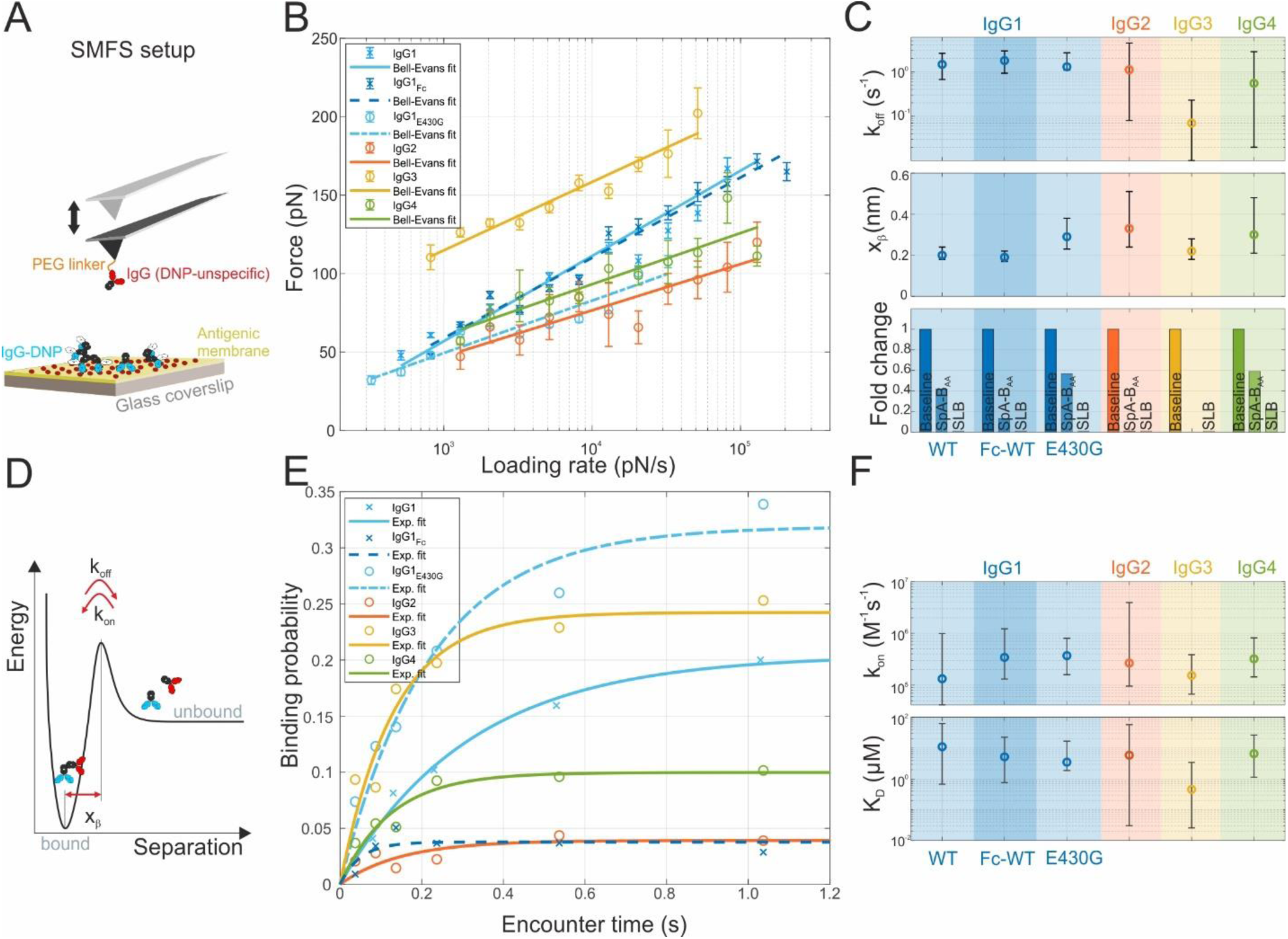
Single molecule force spectroscopy (SMFS) of IgG subclass specific Fc – Fc interactions. (A) Schematic of a SMFS experiment involving a DNP-unspecific IgG covalently linked via a PEG linker to the AFM tip and DNP-specific IgGs bound to a DNP-SLB generated on a glass coverslip. (B) Dependence of the most probable dissociation forces on the force loading rates (symbols) and least squares fits to the Bell-Evans model (solid lines) for each IgG variant. (C) Dissociation rate constant k_off_ (upper panel) and distance from the bound to the transition state xβ (middle panel) obtained from the fits in (B). Specificity controls employing Fc binding SpA-B_AA_ and bare SLBs. (D) Sketch of the Gibbs free energy landscape along the Fc-Fc reaction coordinate (separation). (E) Binding probability (symbols) as a function of the encounter time for each IgG variant. The solid lines are least-squares fits to mono-exponential functions yielding characteristic time constants used to calculate the kinetic on-rate constants. (F) Association rate constants k_on_ (upper panel) obtained from (E) and equilibrium dissociation constants calculated using K_D_ = k_off_/k_on_.

Interestingly, the shape of the force spectra and the kinetic rate constants determined earlier for IgG1 markedly differ from the values determine here, with k_off_ being two orders of magnitude larger, and k_on_ ten times lower than previously observed. We attribute this to the different configurations of the binding partners on the surface and thus different entropic contributions to the binding in the recent vs. the previous experiments, i.e. antigen-bound IgG oligomers with regularly oriented Fc binding sites or attached via flexible polymer linkers mimicking random orientations in solution.

### Determination of kinetic parameters governing the IgG Fab – antigen interaction

In contrast to the short-lived Fc – Fc interactions, the IgG Fab – antigen interactions are of higher affinity and thus accessible via conventional ensemble methods. The lipophilic DNP hapten used as antigen in our model systems is known to partly partition into the hydrophobic region of lipid bilayers modulating the binding of anti-DNP antibodies for different lipid compositions^16,17^.

Since we are employing DNP-SLBs containing different percentages of DNP-conjugated lipids in HS-AFM experiments, we characterized the binding of Fab fragments IgG-Fab-DNP to our DNP-SLBs in grating coupled interferometry (GCI) experiments. Corresponding concentration series of IgG-Fab-DNPs real-time binding and dissociation to/from DNP-SLBs are shown in Figure 3. Analysis of the curves employing a 1:1 Langmuir binding model indeed yielded different kinetic rate constants, in particular significantly different association rate constants *k_on,Fab_* of 3.1 (2.5, 3.9) x 10^4^ M^-1^s^−1^ and 0.6 (0.4, 0.8) x 10^4^ M^-1^s^−1^ for 0.85 % and 5 % DNP-cap-DPPE content in the DNP-SLBs, respectively. These differences are likely a consequence of the abovementioned partly partitioning of the DNP hapten into the hydrophobic regions of the lipid bilayers affecting their collision probabilities with approaching Fab fragments and thus their association rate constants. The difference in dissociation rate constants was less pronounced at 1.3 (1.1, 1.6) x 10^−1^ s^−1^ and 2.1 (1.6, 2.6) x 10^−1^ s^−1^, respectively. Due to material limitations, we did not determine the kinetic rate constants k_on,Fab_ and k_off,Fab_ for the 2.5 % DNP-SLB in GCI experiments, so that for fitting the model to our HS-AFM data k_on,Fab 2.5%_ was adjusted and k_off,Fab 2.5%_ was assumed to be equal to k_off_,_Fab 5%_ (see below).

**Figure 3.**
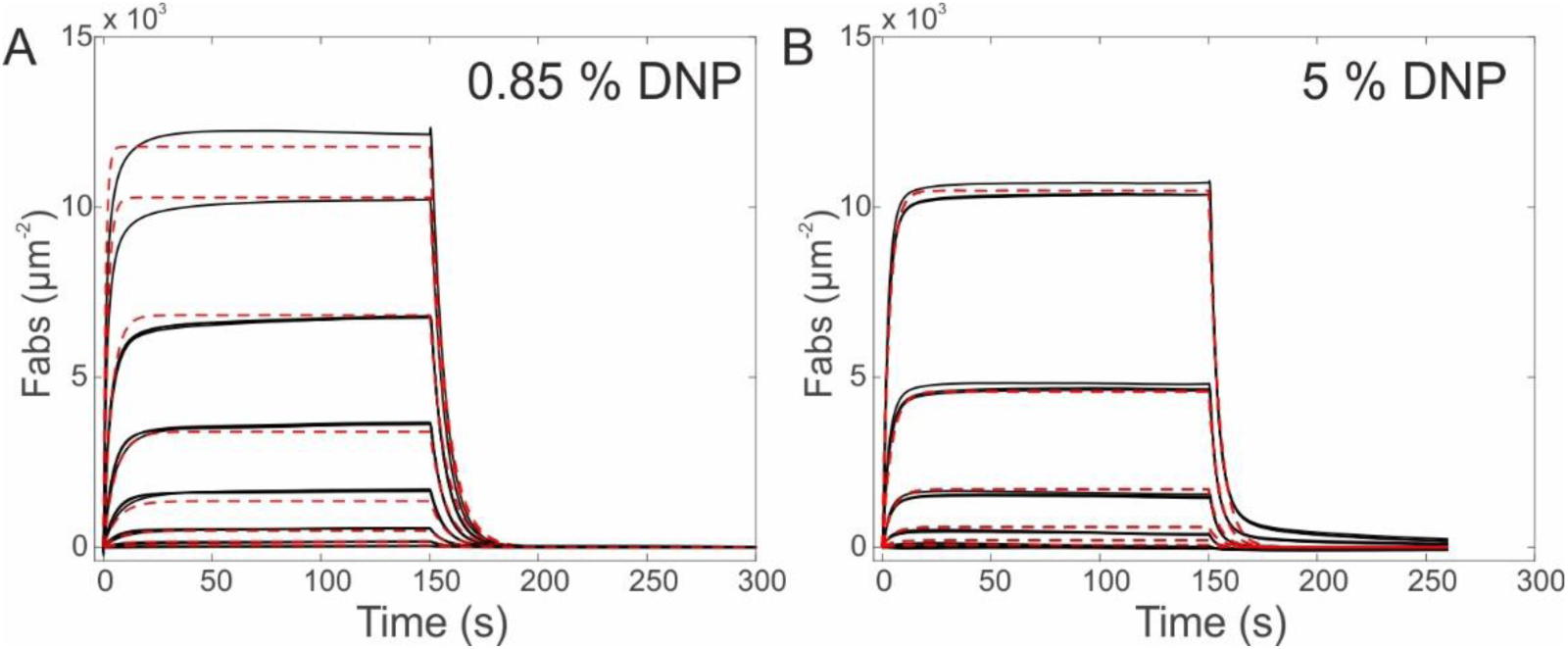
Grating coupled interferometry (GCI) characterization of IgG-Fab-DNP (30 nM, 90 nM, 270 nM, 810 nM, 2.43 µM, 7.29 µM, 14.85 µM) binding to SLBs containing 0.85 %. (A) and 5 % (B) DNP. Dashed lines represent global least-squares fits to a 1:1 Langmuir binding model.

### Fitting the model to IgG1 HS-AFM data

Given that the association rate constants of the IgG-Fab – DNP-cap-DPPE interactions depend on the composition of our DNP-SLBs, all other rates associated with the interaction of the IgG-Fab and DNP (i.e. k_1b_, k_3_ and k_4_) should be similarly affected, which is accounted for by pre-factors α_0.85%_ and α_2.5%_. These pre-factors comprise differences in collision probabilities between the paratope at the tip of the IgG Fab domain and the DNP hapten embedded in the respective SLBs. The rate constants thus modify to k_1b_^i^ = α_i_ k1b, k3^i^ = αi k_3_, and k_4_ ^i^ = α_i_ k_4_, for i = 0.85% and 2.5%. To determine the remaining unknown parameters of our model, we collected IgG oligomer distributions of IgG1-DNP and IgG1-DNP-E430G on DNP-SLBs for 35 different experimental conditions in HS-AFM experiments.

Increasing IgG concentrations or DNP-content of the DNP-SLBs (Figure 4, S3) or increasing IgG incubation times (Figure S4), resulted in enhanced oligomerization, increasing SpA-B_AA_ concentrations during IgG incubation, on the other hand, hampered oligomer formation (Figure S5), grey bars respectively. A global fit of our model (upper branch of the vertical oligomerization pathway, Fig. 1A) to this data (green bars) reasonably reproduces the experimental distributions yielding best fit parameters and corresponding confidence intervals as given in Table 1. The values obtained for the rates k_1b_, k_3,_ and k_4_ are all within the same 10^−4^ s^-1^µm^2^ range reflecting that the respective transitions involve very similar geometries and length scales only slightly differing in flexibility/mobility of the paratope at the end of the unbound Fab tip and the resulting collision probability with an epitope on the DNP-SLB. The E430G point mutation which is associated with an increased flexibility of the IgG Fc domain^4,5,7^ increases these probabilities, resulting in a 1.9-fold (ν_E430G/WT_) increase of rates k_3_ and k_4_. The differences in collision probabilities between IgG Fabs and DNP haptens embedded in different SLBs (cf. Figure 3 and k_on,Fab 2.5% DNP_, Table 1), also impact k_1b_, k_3,_ and k_4_, resulting in 2.0 and 1.5 x higher values (α_0.85%_ and α_2.5%_) for the 0.85 % and 2.5 % DNP-SLBs compared to the 5 % DNP-SLB, respectively. The mass transport rate from the bulk into the surface compartment is in the typical range for such processes^4,10^, and the association rate constant of the Fc – Fc interaction determined from the fit (k_2_) is in the same order of magnitude as the value estimated independently in SMFS experiments (Figure 2).

**Figure 4.**
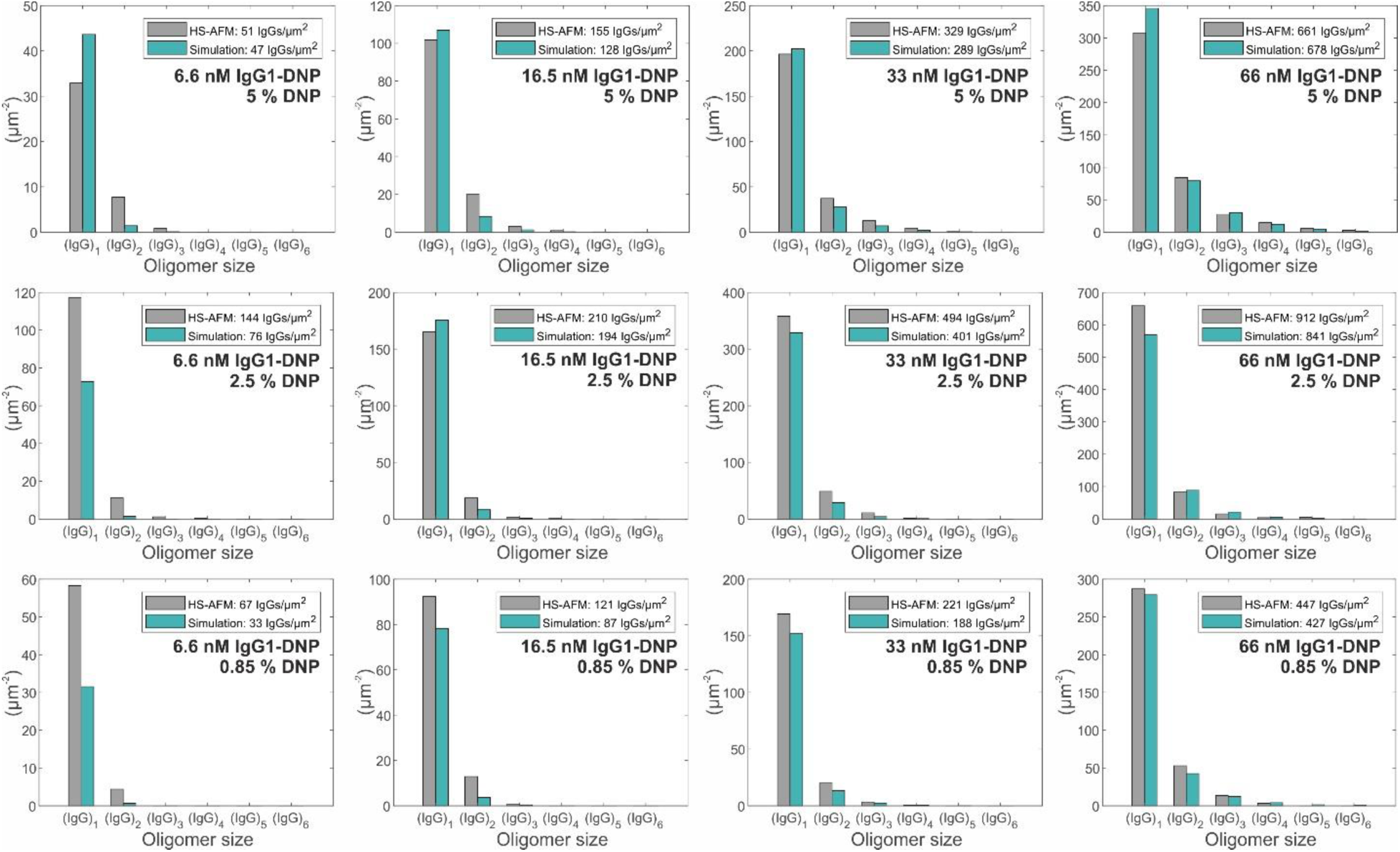
IgG1-DNP oligomer distributions obtained through 3 min incubation of 6.6 – 33 nM IgG1-DNP on SLBs containing 0.85 % DNP-DPPE (lower row), 2.5 % DNP-DPPE (middle row), and 5 % DNP-DPPE (upper row). Grey bars depict experimentally determined oligomer abundances; green bars represent a global fit (together with the data presented in Figure S3-S5) to our kinetic model with parameters given in Table 1. Values in the legends correspond to the total IgG densities.

**Table 1.** Fixed and floating parameters obtained from fitting the mechanistic IgG oligomerization model to HS-AFM IgG1 oligomer distributions.

| **Fixed parameters** | 0.85 % DNP | 2.5 % DNP | 5 % DNP |
| --- | --- | --- | --- |
| $\frac{k_1}{1.7}^{(1)} = k_{on,Fab} (M^{-1}s^{-1})^{(2)}$ | **3.1** x 10<sup>4</sup> | | **0.6** x 10<sup>4</sup> |
| $k_{-1} = k_{off,Fab} (s^{-1})^{(2)}$ | **1.3** x 10<sup>-1</sup> | **2.1** x 10<sup>-1</sup> | **2.1** x 10<sup>-1</sup> |
| $k_{-2} = k_{off,Fc} (s^{-1})^{(3)}$ | **1.5** | | |
| $k_C (M^{-1}s^{-1})^{(4)}$ | **1.9** x 10<sup>5</sup> | | |
| $k_{-C} (s^{-1})^{(4)}$ | **3.2** x 10<sup>-3</sup> | | |
| **Floating parameters** | **best fit (95 % CI)** |  |  |
| $k_{1b} (x 10^{-4} s^{-1}\mu m^2)$ | **3.2** (2.9, 3.6) | | |
| $k_2 (x 10^5 M^{-1}s^{-1})$ | **5.7** (2.8, --) | | |
| $k_3 (x 10^{-4} s^{-1}\mu m^2)$ | **2.5** (0.1, 5.4) | | |
| $k_4 (x 10^{-4} s^{-1}\mu m^2)$ | **6.5** (0.2, 14.4) | | |
| $\alpha_{0.85\%}^{(5)}$ | **2.0** (1.6, 2.3) | | |
| $\alpha_{2.5\%}^{(5)}$ | **1.5** (1.1, 2.2) | | |
| $v_{E430G/WT}^{(6)}$ | **1.9** (1.6, 2.3) | | |
| $k_{on,Fab 2.5\% DNP} (x 10^4 M^{-1}s^{-1})$ | **2.8** (2.3, 3.1) | | |
| $k_T (x 10^{-1} s^{-1})^{(7)}$ | **3.5** (2.7, 4.5) | | |

<sup>(1)</sup> The association rate constant of an IgG for its target epitope is 1.7 x the association rate constant of its Fab fragment<sup>31</sup>. <sup>(2)</sup> Determined from GCI experiments (Figure 3). <sup>(3)</sup> Average of dissociation rate constants determined for IgG1, IgG1-Fc, and IgG1-E430G in SMFS experiments (Figure 2). <sup>(4)</sup> Association and dissociation rate constants of the interaction between IgG1-Fc and SpA-B<sub>AA</sub> taken from<sup>32</sup>. <sup>(5)</sup> Factors modifying rates $k_3$ , $k_4$ , and $k_{1b}$ for 0.85 and 2.5 % DNP-SLBs. <sup>(6)</sup> Factor modifying rates $k_3$ and $k_4$ for the E430G point mutant. <sup>(7)</sup> Mass transport rate from the bulk into the surface compartment is in the typical range for such processes<sup>4,10</sup>.

### Expanding the model to other IgG subclasses

Having determined the model parameters for IgG1 oligomerization we now move on to apply the same strategy to the oligomer distributions obtained for the other IgG subclasses and their E430G point mutants (Figure 5). As found earlier^5^, IgG2 forms oligomers on antigenic membranes so that our model should also be applicable to this IgG subclass. However, as also found in our previous study, it tends to disrupt DNP-SLBs on mica substrates affecting the reproducibility of the collected oligomer distributions, which cannot be accounted for in our model so that we have to limit our analysis to IgG3 and IgG4 subclasses. Sample preparation for these experiments involved a larger droplet volume and as a consequence, the mass transport from the bulk into the surface compartment changed as compared to the previous experiments^18^. We determined the modified rate for this set of experiments from a single parameter fit (k_T_; leaving all other parameters as in Table 1) of the IgG oligomerization model to IgG1 oligomer distributions obtained under identical conditions as for IgG3 and IgG4 (Figure S6). We further set the association rate constants of the Fc – Fc interactions to the value obtained for IgG1 (Table 1), which is justified considering that the respective values obtained via SMFS are not significantly different for all IgG subclasses. A fit of the model to the experimental distributions obtained through adjustment of the remaining free parameters (k_1b_, k_3_, k_4_, and ν_E430G/WT_) is depicted in Figure 5, and the best fit parameters are given in Table 2. For IgG3 and IgG4 subclasses, monovalent-bivalent transition rates k_1b_ have been found to be in the same order of magnitude as for IgG1. The impact of the E430G point mutation on the respective rates (k_3_ and k_4_) was smaller and less significant than for IgG1, in accordance with our previous findings that IgG3 only exhibits marked differences to IgG1 at significantly lower antigen densities^5^. Most strikingly, however, k_3_ was found to be 10-fold lower in case of IgG3 than the corresponding rates for IgG1 and IgG4, which can be rationalized considering that the hinge length of IgG3 is ∼ 4 times longer (62 aa) than the hinges of IgG1 (15 aa) and IgG4 (12 aa). The Fab domains of an IgG3 molecule that is only kept near the antigenic surface via an Fc – Fc interaction (e.g. states y_2,b_, y_4,b_ etc., Figure 1) have a much higher volume to screen for available antigenic epitopes than the shorter-hinge IgGs. While this enables them to reach epitopes further apart from their Fc – Fc anchor as compared to IgG1 and IgG4 thus enabling more efficient oligomerization at low antigen densities, epitope binding (under otherwise identical conditions) will take longer as there is also more space sampled without a potential binding partner. The available time to find an antigenic epitope is limited by the bond lifetime of the Fc – Fc interaction (i.e. 1/k_off,Fc_), which was found to be 8-20 – fold higher in case of IgG3 than for the other subclasses (Figure 2). The combination of increased hinge-length and prolonged Fc – Fc bond lifetime thus enables IgG3 to oligomerize more efficiently than the other subclasses even at low antigen densities^5^. Intriguingly, the IgG3 Fc-Fc interface differs from the other subclasses only by a single aa substitution at position 435 (Arg instead of a His^19^). This, combined with the observation that substitution H435R in an anti-CD20 IgG1 antibody significantly enhanced CDC^1^, suggests that R435 strengthens Fc-Fc interactions in IgG3.

**Figure 5.**
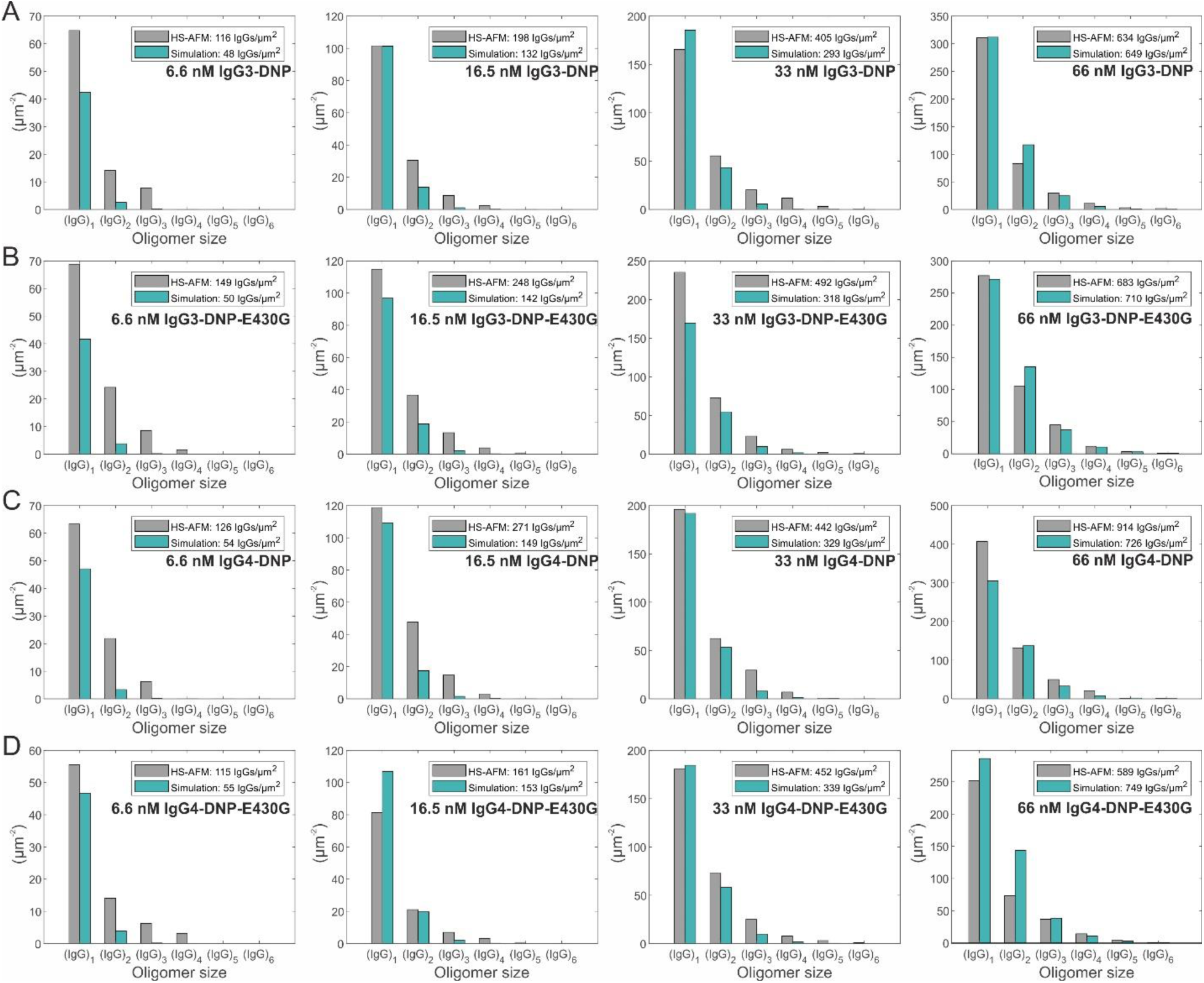
IgG-DNP oligomer distributions obtained through 3 min incubation of 6.6 – 33 nM IgG3-DNP. (A), IgG3-DNP-E430G (B), IgG4-DNP (C), and IgG4-DNP-E430G (D) on SLBs containing 5 % DNP-DPPE. Grey bars depict experimentally determined oligomer abundances; green bars represent a global fit to our kinetic model with parameters given in Table 2.

**Table 2.** Fixed and floating parameters obtained from fitting the mechanistic IgG oligomerization model to HS-AFM IgG3 and IgG4 oligomer distributions.

| Fixed parameters | IgG3 | IgG4 |
| --- | --- | --- |
| $\frac{k_1}{1.7} = k_{on,Fab} (M^{-1}s^{-1})$ <sup>(2)</sup> | **0.6** x 10<sup>4</sup> | |
| $k_{-1} = k_{off,Fab} (s^{-1})$ <sup>(2)</sup> | **2.1** x 10<sup>-1</sup> | |
| $k_2 = k_{on,Fc} (M^{-1}s^{-1})$ <sup>(3)</sup> | **5.7** x 10<sup>5</sup> | |
| $k_{-2} = k_{off,Fc} (s^{-1})$ <sup>(4)</sup> | **0.07** | **0.55** |
| $k_T (x 10^{-1} s^{-1})$ <sup>(5)</sup> | **0.93** (0.7, 1.09) | |
| Floating parameters | best fit (95 % CI) |  |
| $k_{1b} (x 10^{-4} s^{-1}\mu m^2)$ | **1.7** (1.2, 2.4) | **2.2** (1.4, 4.0) |
| $k_3 (x 10^{-4} s^{-1}\mu m^2)$ | **1.2** (0.7, 1.9) x 10<sup>-1</sup> | **1.0** (0.6, 2.0) |
| $k_4 (x 10^{-4} s^{-1}\mu m^2)$ | **2.2** (1.1, 4.0) x 10<sup>-1</sup> | **2.1** (1.1, 4.3) |
| VE430G/WT | **1.2** (0.8, 2.1) | **1.4** (0.8, 2.6) |

<sup>(1)</sup> The association rate constant of an IgG for its target epitope is 1.7 x the association rate constant of its Fab fragment<sup>31</sup>. <sup>(2)</sup> Determined from GCI experiments (Figure 3). <sup>(3)</sup> Taken from the model fit to IgG1 oligomerization, Table 1. <sup>(4)</sup> Dissociation rate constants determined for IgG3 and IgG4 in SMFS experiments (Figure 2). <sup>(5)</sup> Mass transport rate from the bulk into the surface compartment for this set of experiments was determined from a fit of the IgG oligomerization model to IgG1 oligomer distributions obtained under identical conditions as for IgG3 and IgG4 (Figure S6).

### Thermodynamic considerations of IgG oligomerization

Based on the equilibrium dissociation constants 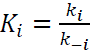 calculated from the kinetic parameters obtained above we can estimate the free energy changes associated with the respective reactions by Δ*G*_*i*_ = *k*_*B*_*T logK*_*i*_, with *k_B_* being Boltzmann’s constant and *T* = 300 K, and assuming unity activity coefficients^11,20^ (Table 3). The decrease of free energy for bivalent binding (ΔG_1b_) is approx. a factor of two larger than for monovalent attachment (ΔG_1_) which is in agreement with earlier studies on similar model systems^11,20^. Assuming equal enthalpies for the two Fab – DNP interactions, entropic differences between monovalent and bivalent binding can be calculated from the respective free energy differences Δ*G*_*i*_ = Δ*H*_*i*_ − *T*Δ*S*_*i*_ (for *i = 1, 1b*) resulting in ΔΔS = Δ*S*_1*b*_ − Δ*S*_1_ = −(Δ*G*_1*b*_ − Δ*G*_1_)/*T*. ΔΔS was found to be 72 to 82 kJ/(mol·K) higher for the second binding event than for the first for all IgGs (Table 3), reflecting that the entropic cost of bringing the antibody to the surface and properly orienting it is almost entirely paid by the first binding event. Subsequent bivalent binding is facilitated by the resulting partial alignment of the unbound Fab towards the antigenic surfaces within the monovalently bound IgG so that ΔS_1b_ mainly contains contributions from lateral diffusion and entropic changes such as the displacement of water molecules from the paratopes / epitopes upon binding^20^. In a similar way, we can characterize entropy differences among complexes along the oligomerization pathways and between different IgG subclass /E430G point mutant variants and relate them to structural features thereof. Again, equal enthalpies for all Fab – DNP interactions can be assumed and, consequently, free energy differences directly relate to the respective entropic contributions to the bindings (Table 3). On a fundamental level, these differences reflect the different numbers of energetically equivalent microstates Ω_*i*_ associated with the respective oligomer configuration (“macrostates” y_i_ as depicted in Figure 2), given by *S*_*i*_ = *k*_*b*_ ln Ω_*i*_. The entropy changes between “macrostates” that involve rates k_3_ or k_4_ (i.e. the transitions from y_2b_ to y_3b_ and y_4b_ to y_5b_ cf. Figure 1 and 6) were found to be higher for the E430G point mutants than for the parental IgGs across subclasses, i.e. Δ*S*_3 *E*430*G*_ > Δ*S*_3_(and Δ*S*_4 *E*430*G*_ > Δ*S*_4_). Expressing this inequality in terms of absolute entropy values reads 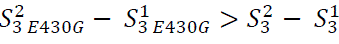, or equivalent 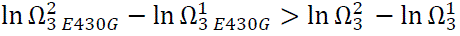 in terms of microstates, with superscript numbers denoting the respective initial (1) and final (2) states of the transitions (Figure 6). In the initial states, 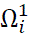 is proportional to the volume V in which the unbound Fabs can screen for a binding partner, whereas in the final states 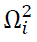 is proportional to the area A on the antigenic membrane within which the Fab can access a potential antigen. The E430G point mutation thus either reduces the volume in which the Fab may search for a binding partner and/or increases the accessible area on the antigenic surfaces. However, given that the mutation increases the flexibility of the Fc domain^7^, the latter scenario is more likely.

**Figure 6.**
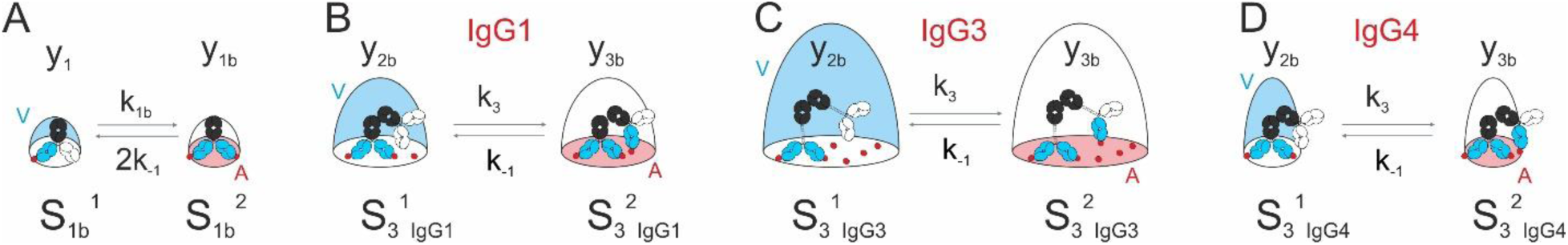
Transitions during IgG oligomerization involving rates k_1b_. (A) and k_3_ (and analogously k_4)_ (B-D). The initial and final macrostates y_i_ each represent a certain number of energetically equivalent microstates Ω_i_ and associated entropy values given by S_i_ = k_B_ ln Ω_i_. The number of microstates in each initial macrostate is proportional to the volume V in which the unbound Fabs can screen for a binding partner. In the respective final macrostates on the other hand, the number of microstates is proportional to the area A on the antigenic membrane within which the Fab can access a potential antigen.

**Table 3.** Thermodynamic parameters deduced from kinetic rate constants.

|  | IgG1 |  |  | IgG3 | IgG4 |
| --- | --- | --- | --- | --- | --- |
|  | 0.85 % DNP | 2.5 % DNP | 5 % DNP | 5 % DNP | 5 % DNP |
| **Gibbs free energy<sup>(1)</sup> $\Delta G$ (kJ/mol)** |  |  |  |  |  |
| $\Delta G_1$ | -32.2 | -30.8 | -26.9 | | |
| $\Delta G_{1b}$ | -54.3 | -52.5 | -51.4 | -49.9 | -50.5 |
| $\Delta G_2$ | -32.1 | | | -39.7 | -34.5 |
| $\Delta G_3$ | -53.7 | -51.9 | -50.8 | -43.3 | -48.6 |
| $\Delta G_4$ | -56.1 | -54.3 | -53.2 | -44.8 | -50.4 |
| $\Delta G_{3, E430G}$ | -55.3 | -53.5 | -52.4 | -43.7 | -49.4 |
| $\Delta G_{4, E430G}$ | -57.7 | -55.9 | -54.8 | -45.2 | -51.2 |
| **Entropy difference<sup>(2)</sup> $\Delta \Delta S$ (kJ/mol K)** |  |  |  |  |  |
| $\Delta S_{1b} - \Delta S_1$ | 73.8 | 72.3 | 81.8 | 76.6 | 78.7 |
| $\Delta S_3 - \Delta S_{3, E430G}$ | -5.3 | | | -1.5 | -2.8 |
| $\Delta S_3 - \Delta S_4$ | -7.9 | | | -5.0 | -6.2 |
| $\Delta S_{3, IgG1} - \Delta S_{3, IgG3/4}$ | | | | 25.2 | 7.6 |

<sup>(1)</sup> Gibbs free energy changes associated with the respective reactions were calculated using $\Delta G_i = k_B T \log K_i$ , with $k_B$ being Boltzmann's constant, absolute temperature $T = 300$ K, equilibrium dissociation constants $K_i = \frac{k_i}{k_{-i}}$ , and assuming unity activity coefficients <sup>11,20</sup>.
<sup>(2)</sup> Entropy differences ( $\Delta \Delta S$ ) were calculated from the respective free energy differences ( $\Delta G_i - \Delta G_j$ ). Using the relation $\Delta G_i = \Delta H_i - T \Delta S_i$ and assuming equal enthalpy ( $\Delta H_i$ ) for all Fab – DNP interactions we get $\Delta \Delta S_{ij} = \Delta S_i - \Delta S_j = -(\Delta G_i - \Delta G_j)/T$ .

Similar considerations can be applied to comparing the transitions associated with k_3_ and k_4_ along the oligomerization pathway of a particular IgG variant which yield Δ*S*_4_ > Δ*S*_3_ (Table 3) and thus 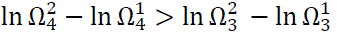. Since the surface area covered by a particular IgG oligomer correlates with the respective area fraction of an IgG hexamer^3^, the area on the antigenic membrane within which the Fab can access a potential antigen is likely comparable in the respective final states. It is thus the number of microstates/ the volume to which the Fab is confined to in state y_2b_ that needs to be larger than in y_4b_, which seems plausible considering that the incoming IgG is bound to a relatively flexible Fc domain of a single bivalently bound antibody in state y_2b_ and can thus screen a larger volume than an IgG that is bound to a partially formed and already aligned Fc platform as in y_4b_.

The entropy difference between monomer-dimer transitions in the IgG1 and IgG3 oligomerization pathways implies that 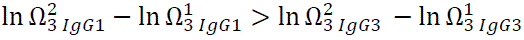. This can again be rationalized considering the accessible areas vs. volumes in the respective macrostates that give rise to a distinct number of associated microstates (Figure 6B vs. 6C). The increased hinge length (l_h_) of IgG3 enables its Fabs to access epitopes within a larger area (proportional to l_h_ ^2^) resulting in an increase of the number of final microstates as compared to 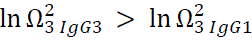. At the same time, however, the associated volume increases more steeply since it is proportional to l_h_^3^, so that 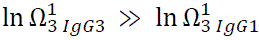 holds thus satisfying Δ*S*_3 *IgG*1_ > Δ*S*_3 *IgG*3_ as determined from the experiment. Comparing IgG1 and IgG4 yields Δ*S*_3 *IgG*1_ > Δ*S*_3 *IgG*4_and thus 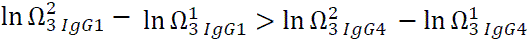. For this inequality to hold, IgG4 either has an increased volume in which the unbound Fab may search for a binding partner and/or a decreased accessible area on the antigenic surfaces, i.e. exactly the opposite effect of the E430G point mutation. However, since IgG4 exhibits a less flexible and shorter hinge than IgG1, a reduction in accessible area seems more plausible than a volume increase (Figure 6B vs. 6C).

### Simulation of complement dependent lysis

Having established a model for IgG oligomerization on antigenic membranes allows us to combine IgG oligomerization and subsequent complement C1 binding to generate a simple kinetic model for simulating liposomal vesicle-based complement lysis assays which may serve as a basis for more complex CDC or PD modelling. We have previously developed a model for the interaction between complement C1 and differently sized IgG oligomers which we successfully applied to real-time QCM data to obtain the kinetic parameters underlying the respective multivalent interactions^5^. Application of this model to IgG oligomer distributions generated from simulating IgG1-DNP and IgG1-DNP-E430G oligomerization for conditions used in DNP-labeled liposomal vesicle-based complement lysis assays^5^ yields surface densities of C1 molecules bound to C1-activating IgG oligomers, i.e. tetramers or larger^3^. To obtain vesicle lysis vs. IgG concentration curves, we assumed these to be Poisson distributed among the vesicles (diameter of 200 nm^21^) and that each active C1 molecule gives rise to the formation of at least one membrane attack complex, which is justified by the fact that vesicles do not possess complement regulatory proteins or other defense mechanisms against complement mediated lysis^22^. Using a lysis threshold of one MAC pore per vesicle, we obtained typical sigmoidal shaped lysis vs. IgG concentration curves (Figure 7A) with EC50 values in the same range as in the corresponding experiment^5^. In addition, the simulation yields the underlying number of active C1 and total IgGs bound per vesicle (Figure 7B and C), the IgG oligomer distributions (Figure 7D) and the corresponding C1 recruitment efficiency (Figure 7E), which are important quantities that are usually not accessible in this type of assay. While the overall outcome of the experiment is thus well predicted, the transition from 0 to 100% lysis extends over a narrower concentration range compared to the experimental curves, where lysis sets on already at lower concentrations. Given that the lipid mixture used in these assays are fluid at room temperature and the DNP molecules thus exhibit a higher lateral mobility than in our gel phase lipid model system on which our simulation is based, the discrepancy is likely a consequence of the lateral oligomerization pathway not being accounted for in our simulations. The lateral pathway is expected to play a more significant role in systems with higher lateral mobility of antigens^3^, especially at lower IgG concentrations where oligomerization through the vertical pathway is generally less efficient (Figure 4). A deviation from the experimental curves could also be a signature of another effect which is not yet considered in the model, i.e. the potential additional IgG recruitment into oligomers (and thus enhancement of oligomerization) via the unbound gC1q heads of IgG-oligomer bound C1 (e.g. the three unbound gC1q heads of an IgG trimer-bound hexameric C1), which could play an important role in both the lateral and vertical oligomerization pathways.

**Figure 7.**
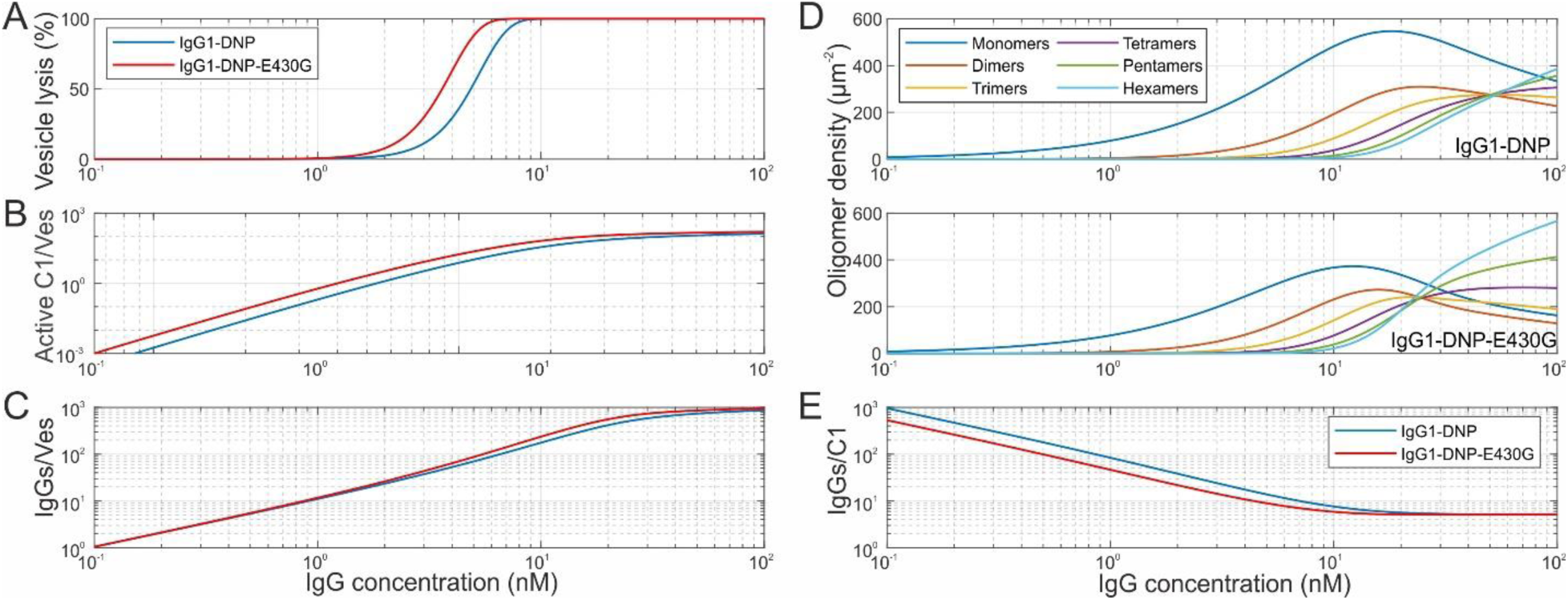
Simulation of liposomal vesicle-based complement lysis assays. (A) Vesicle lysis induced by varying concentrations of IgG1-DNP and IgG1-DNP-E430G. (B) Abundance of C1 bound to IgG1 tetramers or larger per vesicle in dependence of IgG concentration. (C) Total IgGs bound per vesicle. (D) IgG oligomer distributions on the vesicle surfaces in dependence of IgG concentration. (E) C1 recruitment efficiency as a function of the IgG concentration.

## CONCLUSIONS

This study presents a mechanistic model of antigen-dependent IgG oligomerization, that accounts for functionally bivalent IgGs, IgG subclasses, and competitive inhibition of Fc – Fc interactions by SpA-B_AA_. To reduce the number of free parameters in the model, we determined Fab – antigen and Fc – Fc interaction rates for all IgG subclasses in GCI and SMFS experiments, respectively. Interestingly, IgG3 displayed a significantly lower dissociation rate constant compared to IgG1, IgG2, and IgG4, indicating stronger Fc – Fc interactions, which in combination with its increased hinge length enables efficient IgG3 oligomerization even at low antigen densities, where the other subclasses fail to do so. By fitting the model to oligomer distributions obtained from HS-AFM observations of IgG1, IgG3, and IgG4 on antigenic surfaces under various experimental conditions, we have determined the rate constants for crucial steps in the IgG oligomerization process and identified the factors influencing these parameters across IgG subclasses and mutations. Thermodynamic analysis provides insights into the entropic and energetic factors that govern IgG binding to antigenic surfaces and the formation of oligomers. The findings suggest that increased flexibility of the Fc domain (e.g. in the E430G mutant) and hinge length variations (e.g. in IgG3) significantly impact the oligomerization efficiency and thermodynamic stability of the complexes. The upper branch of the vertical oligomerization pathway was sufficient to capture the dynamics observed in our current DNP-SLB model system, which shares similarities—particularly high antigen surface density and low lateral mobility— with bacterial surfaces. Extension of the model to include C1 binding to IgG oligomers further enabled us to generate a simple simulation of complement-dependent vesicle lysis in response to IgG binding that successfully reproduces the overall features of the respective experimental curves (e.g. complement-mediated lysis of DNP-coated liposomes, Ref.^3,5^). These experiments involved fluid-phase lipid vesicles instead of the gel-phase DNP-SLBs used in this work. Resulting discrepancies, particularly at lower IgG concentrations, could be overcome by including the lower branch of the vertical oligomerization pathway (cf. Fig.1B) and lateral oligomerization^4^. Similarly, application of the framework to mammalian cell surfaces (lower antigen surface density and higher lateral mobility), and in particular cancer cells, will require inclusion of the branches/pathways associated with monovalently bound IgGs. This will involve additional parameters (cf. Fig. 1B) that are related via the constraints II-X to the respective parameters of the upper branch of the vertical oligomerization pathway, and rate constants accounting for the lateral diffusion driven association/dissociation of IgG oligomers^4^. A possible additional stabilization of IgG oligomers through C1 binding, which has not yet been taken into account, would also have to be included in such a model. Determination of these parameters will require a new experimental approach for assessing IgG oligomer size, given that HS-AFM is too slow to capture IgG oligomers on fluid-phase lipid bilayers. In this context, we are currently developing concepts involving site-specific labeling of antibodies and/or C1 molecules to make IgG oligomerization on cell surfaces accessible to fluorescence 3D single particle localization microscopy^23^. Further refinement of the model, including these additional mechanisms, will enhance its predictive accuracy and better align the simulations with experimental observations such as CDC assays, and could thus serve as a valuable tool for understanding and optimizing antibody engineering and enhancing antibody-based therapies.

## METHODS

### High-speed atomic force microscopy

HS-AFM (SS-NEX, RIBM Ltd., Ibaraki, JP) was conducted in tapping mode at RT (room temperature, 25°C) in liquid, with typical free amplitudes of 1.5 – 2.5 nm and amplitude setpoints larger than 90%. Silicon nitride cantilevers with electron-beam deposited tips USC-F1.2-k0.15 (NanoWorld® AG, Neuchatel, CH), nominal spring constants of 0.15 N m^−1^, resonance frequencies around 500 kHz, and a quality factor of approx. 2 in liquids were used.

### DNP labeled liposomes

DNP-labeled liposomes consisting of 1,2-dipalmitoyl-sn-glycero-3-phosphocholine (DPPC), 1,2-dipalmitoyl-sn-glycero-3-phosphoethanolamine (DPPE) and 1,2-dipalmitoyl-sn-glycero-3-phosphoethanolamine-N-[6-[(2,4-dinitrophenyl)amino]hexanoyl] (DNP-cap-DPPE) were used to generate supported lipid bilayers (SLBs) on mica and SiO2 substrates. The lipids were purchased from Avanti Polar Lipids, mixed at different ratios of DPPC:DPPE:DNP-cap-DPPE (90:5:5, 90:7.5:2.5, and 90:9.15:0.85, molar ratios), and dissolved in a 2:1 mixture of chloroform and methanol. After the solvents were rotary-evaporated for 30 min, the lipids were again dissolved in chloroform, which was then rotary-evaporated for 30 min. Drying was completed at a high vacuum pump for 2 h. The lipids were dissolved in 500 μL double distilled water while immersed in a water bath at 60°C, flooded with argon, and sonicated for 3 min at 60°C to create small unilamellar vesicles. These were diluted to 2 mg/mL in buffer #1 (10 mM HEPES, 150 mM NaCl, 2 mM CaCl_2_, pH 7.4) and frozen for storage using liquid N_2_. 0.1%, 0.5%, and 5% DNP-cap-DPPE content of SLBs corresponds to 1.6, 8.1 and 81 x 10^3^ DNP molecules/µm^2^.

### Supported lipid bilayers and HS-AFM data evaluation

DNP labeled supported lipid bilayers (DNP-SLBs) for HS-AFM were prepared on muscovite mica. The liposomes were incubated on the freshly cleaved surface (0.5 mg/ml in buffer #1), placed in a humidity chamber to prevent evaporation, and heated to 60°C for 30 min. Then the temperature was gradually reduced to RT within 30 min, followed by exchanging the solution with buffer #1. After 10 min of equilibration at RT, and 15 more buffer exchanges, the SLB was ready for imaging. To passivate any exposed mica, SLBs were incubated with 330 nM IgG1-b12 (isotype control antibody against HIV-1 gp120) ^24^ for 10 min before the molecules of interest were added. The oligomer distributions on DNP-SLBs were analyzed in a two-step process: Individual particle dimensions were determined by HS-AFM, and their oligomeric state was further confirmed via their decay pattern determined in subsequent forced dissociation experiments ^3^. In brief, molecules were scanned in a non-disrupting manner to gauge their number, height, and shape. Subsequently, the scanning force exerted by the HS-AFM cantilever tip is increased (by decreasing the setpoint-amplitude) to dissociate oligomers into their constituent IgGs. Geometric parameters and oligomer decay patterns are combined to assign each IgG assembly its oligomeric state. Data collection per sample/condition was limited to 60 min, all conditions were replicated at least 3 times.

### Construction, expression, and purification of antibody variants

Antibody heavy-chain (HC) expression vectors were constructed by inserting de novo synthesized (Geneart) codon optimized HC coding regions into expression vector pcDNA3.3 (Invitrogen). The HC coding regions consisted of the VH regions of mAbs, CAMPATH (human CD52-specific ^25^), G2a2 (DNP-specific ^26^) or b12 (HIV-1 gp120-specific ^24^), genetically fused to the CH regions of wild-type human IgG1*03, IgG2*01, IgG3*01 or IgG4*01 or one of the mutant variants containing the E430G point mutation ^7^ (EU numbering conventions are used throughout the manuscript). Likewise, separate light-chain expression vectors were constructed by inserting the corresponding VL coding regions in frame with the CL coding regions of the human (J00241) kappa light chain into expression vector pcDNA3.3. All antibodies were produced under serum-free conditions by co-transfecting relevant heavy and light chain expression vectors in FreeStyle™ Expi293F™ cells, using ExpiFectamine™ 293 (LifeTechnologies), according to the manufacturer’s instructions. IgG1, IgG2 and IgG4 antibody variants were purified by protein A affinity chromatography (MabSelect SuRe; GE Health Care), dialyzed overnight to PBS and filter-sterilized over 0.2-µM dead-end filters. Alternatively, IgG3 antibody variants were purified by protein G affinity chromatography (GE Health Care). Purity was determined by CE-SDS and concentration was measured by absorbance at 280 nm (specific extinction coefficients were calculated for each protein). Batches of purified antibody were tested by high-performance size-exclusion chromatography (HP-SEC) for aggregates or degradation products and shown to be at least 95% monomeric. Purified antibodies were stored at 2 – 8°C.

### Single molecule force spectroscopy

SMFS measurements were conducted on a JPK NanoWizard 4 (Bruker Corporation, MA, US). Measurements were performed in buffer #1 at pH 7.4. Commercial cantilever chips with nominal spring constants of 10 – 20 pN/nm (model MSCT, Bruker, USA) were used. Sensitivity and spring constant calibration of the cantilever were done using the thermal noise method and the JPK calibration manager. Functionalization of AFM cantilevers with DNP-unspecific IgGs (IgG-b12, IgG-CD52) and mutant variants was performed as described^27^. For the determination of the dissociation rate constants, FD curves were recorded at 6 different pulling rates between 200 nm/s and 20000 nm/s with each dataset containing 1000 FD curves. The total encounter time was kept constant between loading rates. Unbinding forces were binned with respect to the loading rate and the resulting force spectra were analyzed employing the Bell-Evans model^12^. Data analysis for SMFS was performed using a MATLAB (Mathworks) routine developed in-house. Specificity controls were done by introducing 4.5 µM SpA-B_AA_^9^ into the AFM liquid cell and evaluating the resulting reduction of binding probability and additionally by recording FD curves on SLBs before the incubation with DNP-specific IgGs.

Association rate constants k_on,Fc_ were determined as described earlier^28^. Briefly, six encounter times between 0 ms and 1000 ms were chosen, and the resulting binding probability p_b_ (measured at a pulling velocity 1000 nm/s) was determined from 1000 curves at each setting. Plots of p_b_ over encounter time were fitted by 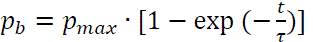, with p_max_ being the maximum observable binding probability, t being the encounter time, and τ being the characteristic time constant of association. Estimation of k_on_ according to k_on_=(τ·c_eff_)^−1^ requires knowledge of the effective concentration c_eff_ which describes the number of binding partners (IgGs) within the effective volume V_eff_ accessible for free equilibrium interaction. V_eff_ can be described as a half-sphere with radius r_eff_, which is given by the average unbinding length determined from the force vs. distance curves. For the interaction of two IgGs there are 4 different realizations of Fc-Fc interactions (both IgGs have two Fc-Fc interfaces) so that c_eff_ finally reads 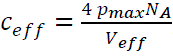, with N_A_ being Avogadro’s constant. For each setting in a square spot of 500 nm at least 1000 FD cycles evenly distributed over a 10 x 10 grid were recorded. Measurements were performed in buffer #1.

### Grating coupled interferometry

Grating-coupled interferometry (GCI) experiments were conducted to obtain kinetic data for IgG1-Fab with the Creoptix WAVE delta (Malvern Panalytical, Wädenswill, Switzerland). The 4PCP-LIP WAVE chip (pre-immobilized lipophilic groups for capturing lipid-based particles or hydrophobic ligands, Creoptix) was primed with borate buffer (1 M borate, 1 M NaCl, pH 9.0) and PBS. Following the priming procedure, the running buffer was replaced with buffer #1 and the flow rate was set to 40 µL/min per channel. Unless otherwise stated, all subsequent steps were conducted under these conditions. The probes were stored in the autosampler during the experiment at a temperature of 6 °C. The conditioning of the chip was performed using three cycles of CHAPS (20nM), with each cycle lasting 120 seconds followed by a running buffer phase. Following the three start-up cycles, DNP labeled liposomes (2 mg/mL) containing varying concentrations of DNP (0%, 0.85%, and 5%) was incubated for 1200 seconds with a flow rate of 2.5 µL/min per channel on the chip. One of the four channels was designated as the baseline, and the lipid without DNP was incubated in that channel. Following an additional flushing step with CB, the lipid membrane was passivated two times with NaOH (10 mM), then a non-binding control antibody IgG1-b12 (50 µg/mL) was incubated to check the consistent of the lipid. After a 30-minute running buffer phase, IgG1-Fab was incubated in rising concentrations (30 nM, 90 nM, 270 nM, 810 nM, 2.43 µM, 7.29 µM, 14.85 µM), commencing with a blank cycle (running buffer without IgG1-Fab) for 150 seconds. Each of the IgG Fab incubations was interspersed with a dissociation phase in running buffer for a period of 1500 seconds between the incubations.

### Data fitting and simulations

Fitting of IgG oligomerization models to HS-AFM data (Figure 1 and S1) and simulation of vesicle lysis assays was done in MATLAB (The MathWorks Inc., MA, US) using the ODE15s and fminsearch nonlinear solver (for data fitting). Simulation of IgG oligomer distributions involved the following steps to closely resemble the experimental procedure: Incubation of a certain IgG concentration for a defined time was followed by a washing step (3 min duration) from which on the protein concentration in the bulk compartment was set to 0. After the washing step, IgG oligomer densities were simulated until t = 60 min (corresponding to the experimental data collection duration) and time averaged over this period. Confidence intervals of fit parameters were determined according to^29,30^.

## ASSOCIATED CONTENT

### Supporting Information

The Supporting Information is available free of charge.

Rate equations accounting for the presence of an IgG Fc-Fc interactions inhibiting protein (SpA-B_AA_) and corresponding scheme, additional IgG3 SMFS data, additional experimental IgG1-DNP / E430G oligomer distributions and model fits (PDF).

## AUTHOR INFORMATION

### Corresponding Author

**Johannes Preiner** – University of Applied Sciences Upper Austria, Linz, Austria;

### Other Authors

Jürgen Strasser – University of Applied Sciences Upper Austria, Linz Nikolaus Frischauf – University of Applied Sciences Upper Austria, Linz Lukas Schustereder – University of Applied Sciences Upper Austria, Linz Andreas Karner – University of Applied Sciences Upper Austria, Linz Frank J. Beurskens – Genmab Utrecht, The Netherlands Sieto Bosgra – Genmab Utrecht, The Netherlands Aran F. Labrijn – Genmab Utrecht, The Netherlands

### Author Contributions

The manuscript was written through contributions of all authors. All authors have given approval to the final version of the manuscript. J.S., A.F.L, F.J.B and J.P. designed research. N.F., J.S., L.S., A.K. performed experiments and analyzed data. S.B. reviewed the model. J.P. supervised the project, developed the model, did data analysis, prepared the figures.

### Funding Sources

J.P. acknowledges support the Austrian Science Fund (FWF, Grant No. P33958 and P34164) and the Federal State of Upper Austria as a part of the FH Upper Austria Center of Excellence for Technological Innovation in Medicine (TIMed Center).

### Notes

The authors declare the following competing financial interest(s): F.J.B., S.B., and A.F.L., are inventors on patent applications related to complement activation by therapeutic antibodies and own Genmab stock. J.P. received Genmab funding.

## ACKNOWLEDGMENT

We thank Prof. S. Rooijakkers for providing SpA-B_AA_. This research was funded in whole or in part by the Austrian Science Fund (FWF) [Grant No. P33958 and P34164 to J.P.]. For open access purposes, the author has applied a CC BY public copyright license to any author accepted manuscript version arising from this submission. J.P. acknowledges support by the Federal State of Upper Austria as a part of the FH Upper Austria Center of Excellence for Technological Innovation in Medicine (TIMed Center).

## ABBREVIATIONS

IgG immunoglobulin, CCP classical complement pathway, MAC membrane attack complex, CDC complement dependent cytotoxicity, ADCC antibody-dependent cellular cytotoxicity, ADCP antibody-dependent cellular phagocytosis, SLB supported lipid bilayer, SpA-B_AA_ B domain of staphylococcal protein A, HS-AFM high-speed atomic force microscopy, QCM quartz crystal microbalance, GCI grating coupled interferometry, SMFS single molecule force spectroscopy

