## Supplementary Text and Figures for "A kinetic model of antigen-dependent IgG oligomerization and complement binding"

KEYWORDS

SI TEXT

**Incorporation of a Fc-Fc interactions inhibiting protein into the model.** In the presence of an IgG Fc-domain and thus Fc-Fc interactions inhibiting protein such as staphylococcal protein A (SpA-B)[1], the mechanistic model modifies as outlined in Fig. S1. While only a simple monovalent interaction (i.e. the SpA-B - IgG Fc interaction characterized through the dissociation and association rate constants $k_C$ and $k_{-C}$) is added to the system, the scheme becomes markedly more complex since every IgG oligomer state of the initial model now exists in four different SpA-B - bound states, i.e. with either 0, 1 (two different configurations), or 2 SpA-B bound. Additionally, IgGs in solution (bulk and surface compartment) can also bind SpA-B which must be taken into account. The corresponding rate equations for binding of SpA-B to IgGs in bulk solution thus read (cf. Fig. S1)

$$\frac{dy_{01}}{dt} = -\left(2k_C y_{01} y_{04} - k_{-C} y_{02}\right)$$

$$\frac{dy_{02}}{dt} = \left(2k_C y_{01} y_{04} - k_{-C} y_{02}\right) - \left(k_C y_{02} y_{04} - 2k_{-C} y_{03}\right)$$

$$\frac{dy_{03}}{dt} = \left(k_C y_{02} y_{04} - 2k_{-C} y_{03}\right)$$

$$\frac{dy_{04}}{dt} = -\left(2k_C y_{01} y_{04} - k_{-C} y_{02}\right) - \left(k_C y_{02} y_{04} - 2k_{-C} y_{03}\right)$$

Given that the volume of the liquid droplet applied to DNP-SLBs (~2 µl) is much larger than the volume of the surface compartment (mica substrate diameter of 1.5 mm x 1 µm surface compartment height ≈ 2 pl) we assume constant concentrations $y_{01} - y_{04}$ as soon as equilibrium is established in solution. The rate equations for antigen-bound IgGs are given by

$$\frac{dy_1}{dt} = k_T(y_{01} - y_1) - 2k_C y_{58} y_1 + k_{-C} y_2 - \alpha\left[k_1 y_1 y_{57} - k_{-1} y_4 + 2k_2 y_1(2y_7 + y_8) - k_{-2}(y_{10} + y_{11}) + \sum_{i=14,22,30,38} 2k_2 y_1(y_i + y_{i+1}) - k_{-2}(y_{i+4} + y_{i+5})\right]$$

$$\frac{dy_2}{dt} = k_T(y_{02} - y_2) + k_C y_{58}(2y_1 - y_2) - k_{-C}(y_2 - 2y_3) - \alpha\left[k_1 y_2 y_{57} - k_{-1} y_5 + \sum_{i=7,14,22,33,38} k_2 y_2(2y_i + y_{i+1}) - \sum_{i=12,20,28,36,44} k_{-2} y_2(y_i + y_{i+1})\right]$$

$$\frac{dy_3}{dt} = k_T(y_{03} - y_3) + k_C y_{58} y_2 - 2k_{-C} y_3 - \alpha[k_1 y_3 y_{57} - k_{-1} y_6]$$

$$\frac{dy_4}{dt} = (k_1 y_1 y_{57} - k_{-1} y_4) - (2k_C y_{58} y_4 - k_{-C} y_5) - (k_{1b} y_4 y_{57} - 2k_{-1} y_7)$$

$$\frac{dy_5}{dt} = (k_1 y_2 y_{57} - k_{-1} y_5) - (k_C y_{58} y_5 - 2k_{-C} y_6) - (k_{1b} y_5 y_{57} - 2k_{-1} y_8) + (2k_C y_{58} y_4 - k_{-C} y_5)$$

$$\frac{dy_6}{dt} = (k_1 y_3 y_{57} - k_{-1} y_6) - (k_{1b} y_6 y_{57} - 2k_{-1} y_9) + (k_C y_{58} y_5 - 2k_{-C} y_6)$$

$$\frac{dy_7}{dt} = (k_{1b} y_4 y_{57} - 2k_{-1} y_7) - (2k_C y_{58} y_7 - k_{-C} y_8) - (4k_2 y_7 y_1 - k_{-2} y_{10}) - (2k_2 y_7 y_2 - k_{-2} y_{13})$$

$$\frac{dy_8}{dt} = (k_{1b} y_5 y_{57} - 2k_{-1} y_8) - (k_C y_{58} y_8 - 2k_{-C} y_9) - (k_2 y_8 y_2 - k_{-2} y_{12}) - (2k_2 y_8 y_1 - k_{-2} y_{11}) + (2k_C y_{58} y_7 - k_{-C} y_8)$$

$$\frac{dy_9}{dt} = (k_{1b} y_6 y_{57} - 2k_{-1} y_9) + (k_C y_{58} y_8 - 2k_{-C} y_9)$$

$$\frac{dy_{10}}{dt} = (4k_2 y_7 y_1 - k_{-2} y_{10}) - (k_C y_{58} y_{10} - k_{-C} y_{11}) - (k_3 y_{10} y_{57} - k_{-1} y_{14}) - (k_C y_{58} y_{10} - k_{-C} y_{13})$$

$$\frac{dy_{11}}{dt} = (2k_2 y_8 y_1 - k_{-2} y_{11}) - (k_C y_{58} y_{11} - k_{-C} y_{12}) - (k_3 y_{11} y_{57} - k_{-1} y_{15}) + (k_C y_{58} y_{10} - k_{-C} y_{11})$$

$$\frac{dy_{12}}{dt} = (k_2 y_8 y_2 - k_{-2} y_{12}) + (k_C y_{58} y_{13} - k_{-C} y_{12}) - (k_3 y_{12} y_{57} - k_{-1} y_{16}) + (k_C y_{58} y_{11} - k_{-C} y_{12})$$

$$\frac{dy_{13}}{dt} = (2k_2 y_7 y_2 - k_{-2} y_{13}) + (k_C y_{58} y_{10} - k_{-C} y_{13}) - (k_3 y_{13} y_{57} - k_{-1} y_{17}) - (k_C y_{58} y_{13} - k_{-C} y_{12})$$

$$\frac{dy_{14}}{dt} = (k_3 y_{10} y_{57} - k_{-1} y_{14}) - (k_C y_{58} y_{14} - k_{-C} y_{15}) - (2k_2 y_{14} y_1 - k_{-2} y_{18}) - (2k_2 y_{14} y_2 - k_{-2} y_{21}) - (k_C y_{58} y_{14} - k_{-C} y_{17})$$

$$\frac{dy_{15}}{dt} = (k_3 y_{11} y_{57} - k_{-1} y_{15}) - (k_C y_{58} y_{15} - k_{-C} y_{16}) - (k_2 y_{15} y_2 - k_{-2} y_{20}) - (2k_2 y_{15} y_1 - k_{-2} y_{19}) + (k_C y_{58} y_{14} - k_{-C} y_{15})$$

$$\frac{dy_{16}}{dt} = (k_3 y_{12} y_{57} - k_{-1} y_{16}) + (k_C y_{58} y_{17} - k_{-C} y_{16}) + (k_C y_{58} y_{15} - k_{-C} y_{16})$$

$$\frac{dy_{17}}{dt} = (k_3 y_{13} y_{57} - k_{-1} y_{17}) + (k_C y_{58} y_{14} - k_{-C} y_{17}) - (k_C y_{58} y_{17} - k_{-C} y_{16})$$

$$\frac{dy_{18}}{dt} = (2k_2 y_{14} y_1 - k_{-2} y_{18}) - (k_C y_{58} y_{18} - k_{-C} y_{19}) - (k_4 y_{18} y_{57} - k_{-1} y_{22}) - (k_C y_{58} y_{18} - k_{-C} y_{21})$$

$$\frac{dy_{19}}{dt} = (2k_2 y_{15} y_1 - k_{-2} y_{19}) - (k_C y_{58} y_{19} - k_{-C} y_{20}) - (k_4 y_{19} y_{57} - k_{-1} y_{23}) + (k_C y_{58} y_{18} - k_{-C} y_{19})$$

$$\frac{dy_{20}}{dt} = (k_2 y_{15} y_2 - k_{-2} y_{20}) + (k_C y_{58} y_{21} - k_{-C} y_{20}) - (k_4 y_{20} y_{57} - k_{-1} y_{24}) + (k_C y_{58} y_{19} - k_{-C} y_{20})$$

$$\frac{dy_{21}}{dt} = (2k_2 y_{14} y_2 - k_{-2} y_{21}) + (k_C y_{58} y_{18} - k_{-C} y_{21}) - (k_4 y_{21} y_{57} - k_{-1} y_{25}) - (k_C y_{58} y_{21} - k_{-C} y_{20})$$

$$\frac{dy_{22}}{dt} = (k_4 y_{18} y_{57} - k_{-1} y_{22}) - (k_C y_{58} y_{22} - k_{-C} y_{23}) - (2k_2 y_{22} y_1 - k_{-2} y_{26}) - (2k_2 y_{22} y_2 - k_{-2} y_{29}) - (k_C y_{58} y_{22} - k_{-C} y_{25})$$

$$\frac{\mathrm{d}y_{23}}{\mathrm{d}t} = (\mathrm{k}_4 y_{19} y_{57} - \mathrm{k}_{-1} y_{23}) - (\mathrm{k}_C y_{58} y_{23} - \mathrm{k}_{-C} y_{24}) - (\mathrm{k}_2 y_{23} y_2 - \mathrm{k}_{-2} y_{28}) - (2\mathrm{k}_2 y_{23} y_1 - \mathrm{k}_{-2} y_{27}) + (\mathrm{k}_C y_{58} y_{22} - \mathrm{k}_{-C} y_{23})$$

$$\frac{dy_{24}}{dt} = (k_4 y_{20} y_{57} - k_{-1} y_{24}) + (k_C y_{58} y_{25} - k_{-C} y_{24}) + (k_C y_{58} y_{23} - k_{-C} y_{24})$$

$$\frac{dy_{25}}{dt} = (k_4 y_{21} y_{57} - k_{-1} y_{25}) + (k_C y_{58} y_{22} - k_{-C} y_{25}) - (k_C y_{58} y_{25} - k_{-C} y_{24})$$

$$\frac{dy_{26}}{dt} = (2k_2 y_{22} y_1 - k_{-2} y_{26}) - (k_C y_{58} y_{26} - k_{-C} y_{27}) - (k_5 y_{26} y_{57} - k_{-1} y_{30}) - (k_C y_{58} y_{26} - k_{-C} y_{29})$$

$$\frac{\mathrm{d}y_{27}}{\mathrm{d}t} = (2\mathrm{k}_2 y_{23} y_1 - \mathrm{k}_{-2} y_{27}) - (\mathrm{k}_C y_{58} y_{27} - \mathrm{k}_C y_{28}) - (\mathrm{k}_5 y_{27} y_{57} - \mathrm{k}_{-1} y_{31}) + (\mathrm{k}_C y_{58} y_{26} - \mathrm{k}_{-C} y_{27})$$

$$\frac{dy_{28}}{dt} = (k_2 y_{23} y_2 - k_{-2} y_{28}) + (k_C y_{58} y_{29} - k_{-C} y_{28}) - (k_5 y_{28} y_{57} - k_{-1} y_{32}) + (k_C y_{58} y_{27} - k_{-C} y_{28})$$

$$\frac{dy_{29}}{dt} = (2k_2 y_{22} y_2 - k_{-2} y_{29}) + (k_C y_{58} y_{26} - k_{-C} y_{29}) - (k_5 y_{29} y_{57} - k_{-1} y_{33}) - (k_C y_{58} y_{29} - k_{-C} y_{28})$$

$$\frac{dy_{30}}{dt} = (k_4 y_{26} y_{57} - k_{-1} y_{30}) - (k_C y_{58} y_{30} - k_{-C} y_{31}) - (2k_2 y_{30} y_1 - k_{-2} y_{34}) - (2k_2 y_{30} y_2 - k_{-2} y_{37}) - (k_C y_{58} y_{30} - k_{-C} y_{33})$$

$$\frac{dy_{31}}{dt} = (k_4 y_{27} y_{57} - k_{-1} y_{31}) - (k_C y_{58} y_{31} - k_{-C} y_{32}) - (k_2 y_{31} y_2 - k_{-2} y_{36}) - (2k_2 y_{31} y_1 - k_{-2} y_{35}) + (k_C y_{58} y_{30} - k_{-C} y_{31})$$

$$\frac{dy_{32}}{dt} = (k_4 y_{28} y_{57} - k_{-1} y_{32}) + (k_C y_{58} y_{33} - k_{-C} y_{32}) + (k_C y_{58} y_{31} - k_{-C} y_{32})$$

$$\frac{dy_{33}}{dt} = (k_4 y_{29} y_{57} - k_{-1} y_{33}) + (k_C y_{58} y_{30} - k_{-C} y_{33}) - (k_C y_{58} y_{33} - k_{-C} y_{32})$$

$$\frac{dy_{34}}{dt} = (2k_2 y_{30} y_1 - k_{-2} y_{34}) - (k_C y_{58} y_{34} - k_{-C} y_{35}) - (k_4 y_{34} y_{57} - k_{-1} y_{38}) - (k_C y_{58} y_{34} - k_{-C} y_{37})$$

$$\frac{dy_{35}}{dt} = (2k_2 y_{31} y_1 - k_{-2} y_{35}) - (k_C y_{58} y_{35} - k_{-C} y_{36}) - (k_4 y_{35} y_{57} - k_{-1} y_{39}) + (k_C y_{58} y_{34} - k_{-C} y_{35})$$

$$\frac{dy_{36}}{dt} = (k_2 y_{31} y_2 - k_{-2} y_{36}) + (k_C y_{58} y_{37} - k_{-C} y_{36}) - (k_4 y_{36} y_{57} - k_{-1} y_{40}) + (k_C y_{58} y_{35} - k_{-C} y_{36})$$

$$\frac{dy_{37}}{dt} = (2k_2 y_{30} y_2 - k_{-2} y_{37}) + (k_C y_{58} y_{34} - k_{-C} y_{37}) - (k_4 y_{37} y_{57} - k_{-1} y_{41}) - (k_C y_{58} y_{37} - k_{-C} y_{36})$$

$$\frac{dy_{38}}{dt} = (k_4 y_{34} y_{57} - k_{-1} y_{38}) - (k_C y_{58} y_{38} - k_{-C} y_{39}) - (2k_2 y_{38} y_1 - k_{-2} y_{42}) - (2k_2 y_{38} y_2 - k_{-2} y_{45}) - (k_C y_{58} y_{38} - k_{-C} y_{41})$$

$$\frac{dy_{39}}{dt} = (k_4 y_{35} y_{57} - k_{-1} y_{39}) - (k_C y_{58} y_{39} - k_{-C} y_{40}) - (k_2 y_{39} y_2 - k_{-2} y_{44}) - (2k_2 y_{39} y_1 - k_{-2} y_{43}) + (k_C y_{58} y_{38} - k_{-C} y_{39})$$

$$\frac{dy_{40}}{dt} = (k_4 y_{36} y_{57} - k_{-1} y_{40}) + (k_C y_{58} y_{41} - k_{-C} y_{40}) + (k_C y_{58} y_{39} - k_{-C} y_{40})$$

$$\frac{dy_{41}}{dt} = (k_4 y_{37} y_{57} - k_{-1} y_{41}) + (k_C y_{58} y_{38} - k_{-C} y_{41}) - (k_C y_{58} y_{41} - k_{-C} y_{40})$$

$$\frac{dy_{42}}{dt} = (2k_2 y_{38} y_1 - k_{-2} y_{42}) - (k_C y_{58} y_{42} - k_{-C} y_{43}) - (k_4 y_{42} y_{57} - k_{-1} y_{46}) - (k_C y_{58} y_{42} - k_{-C} y_{45}) + (k_{1b} 5 y_{54} y_{57} - k_{-1} y_{42})$$

$$\frac{dy_{43}}{dt} = (2k_2 y_{39} y_1 - k_{-2} y_{43}) - (k_C y_{58} y_{43} - k_{-C} y_{44}) - (k_4 y_{43} y_{57} - k_{-1} y_{47}) + (k_C y_{58} y_{42} - k_{-C} y_{43})$$

$$\frac{dy_{44}}{dt} = (k_2 y_{39} y_2 - k_{-2} y_{44}) + (k_C y_{58} y_{45} - k_{-C} y_{44}) - (k_4 y_{44} y_{57} - k_{-1} y_{48}) + (k_C y_{58} y_{43} - k_{-C} y_{44})$$

$$\frac{dy_{45}}{dt} = (2k_2 y_{38} y_2 - k_{-2} y_{45}) + (k_C y_{58} y_{42} - k_{-C} y_{45}) - (k_4 y_{45} y_{57} - k_{-1} y_{49}) - (k_C y_{58} y_{45} - k_{-C} y_{44})$$

$$\frac{dy_{46}}{dt} = (k_4 y_{42} y_{57} - k_{-1} y_{46}) - (k_C y_{58} y_{46} - k_{-C} y_{47}) + (k_{1b} 6 y_{50} y_{57} - k_{-1} y_{46}) - (k_C y_{58} y_{46} - k_{-C} y_{49})$$

$$\frac{dy_{47}}{dt} = (k_4 y_{43} y_{57} - k_{-1} y_{47}) - (k_C y_{58} y_{47} - k_{-C} y_{48}) + (k_{1b} 6 y_{51} y_{57} - k_{-1} y_{47}) + (k_C y_{58} y_{46} - k_{-C} y_{47})$$

$$\frac{dy_{48}}{dt} = (k_4 y_{44} y_{57} - k_{-1} y_{48}) + (k_C y_{58} y_{49} - k_{-C} y_{48}) + (k_{1b} 6 y_{52} y_{57} - k_{-1} y_{48}) + (k_C y_{58} y_{47} - k_{-C} y_{48})$$

$$\frac{dy_{49}}{dt} = (k_4 y_{45} y_{57} - k_{-1} y_{49}) + (k_C y_{58} y_{46} - k_{-C} y_{49}) + (k_{1b} 6 y_{53} y_{57} - k_{-1} y_{49}) - (k_C y_{58} y_{49} - k_{-C} y_{48})$$

$$\frac{dy_{50}}{dt} = -(k_{1b} 6 y_{50} y_{57} - k_{-1} y_{46}) - (k_C y_{58} y_{50} - k_{-C} y_{51}) - (k_6 y_{50} - 6 k_{-2} y_{56}) - (k_C y_{58} y_{50} - k_{-C} y_{53}) + (k_4 y_{54} y_{57} - k_{-1} y_{50})$$

$$\frac{dy_{51}}{dt} = -(k_{1b} 6 y_{51} y_{57} - k_{-1} y_{47}) - (k_C y_{58} y_{51} - k_{-C} y_{52}) + (k_C y_{58} y_{50} - k_{-C} y_{51})$$

$$\frac{dy_{52}}{dt} = -(k_{1b} 6 y_{52} y_{57} - k_{-1} y_{48}) + (k_C y_{58} y_{53} - k_{-C} y_{52}) + (k_C y_{58} y_{51} - k_{-C} y_{52})$$

$$\frac{dy_{53}}{dt} = -(k_{1b} 6 y_{53} y_{57} - k_{-1} y_{49}) + (k_C y_{58} y_{50} - k_{-C} y_{53}) - (k_C y_{58} y_{53} - k_{-C} y_{52})$$

$$\frac{dy_{54}}{dt} = -(k_{1b} 5 y_{54} y_{57} - k_{-1} y_{42}) - (k_4 y_{54} y_{57} - k_{-1} y_{50}) - (k_5 y_{54} - k_{-2} y_{55})$$

$$\frac{dy_{55}}{dt} = (k_5 y_{54} - k_{-2} y_{55}) - (k_7 y_{55} - 6 k_{-1} y_{56})$$

$$\frac{dy_{56}}{dt} = (k_6 y_{50} - 6 k_{-2} y_{56}) + (k_7 y_{55} - 6 k_{-1} y_{56})$$

$$\frac{dy_{57}}{dt} = -\sum_{i=4}^{6} \frac{dy_i}{dt} w_{1,m} - \sum_{i=7}^{13} \frac{dy_i}{dt} w_1 - \sum_{i=14}^{21} \frac{dy_i}{dt} w_2 - \sum_{i=22}^{29} \frac{dy_i}{dt} w_3 - \sum_{i=30}^{37} \frac{dy_i}{dt} w_4 - \sum_{i=38}^{45} \frac{dy_i}{dt} w_5 - \sum_{i=46}^{56} \frac{dy_i}{dt} w_6$$

$$\frac{dy_{58}}{dt} = k_T (y_{62} - y_{58}) - k_C y_{58} (2y_1 + y_2) + k_{-C} (y_2 + 2y_3) - \alpha \Bigg[ k_C y_{58} (2y_4 + y_5 + 2y_7 + y_8) - k_{-C} (y_5 + 2y_6 + y_8 + 2y_9) + \sum_{i=0}^{10} 2 k_c y_{58} y_{4i+10} + (k_c y_{58} - k_{-C}) \cdot (y_{4i+11} + y_{4i+13}) - 2 k_{-C} y_{4i+12} \Bigg]$$

### SUPPORTING FIGURES

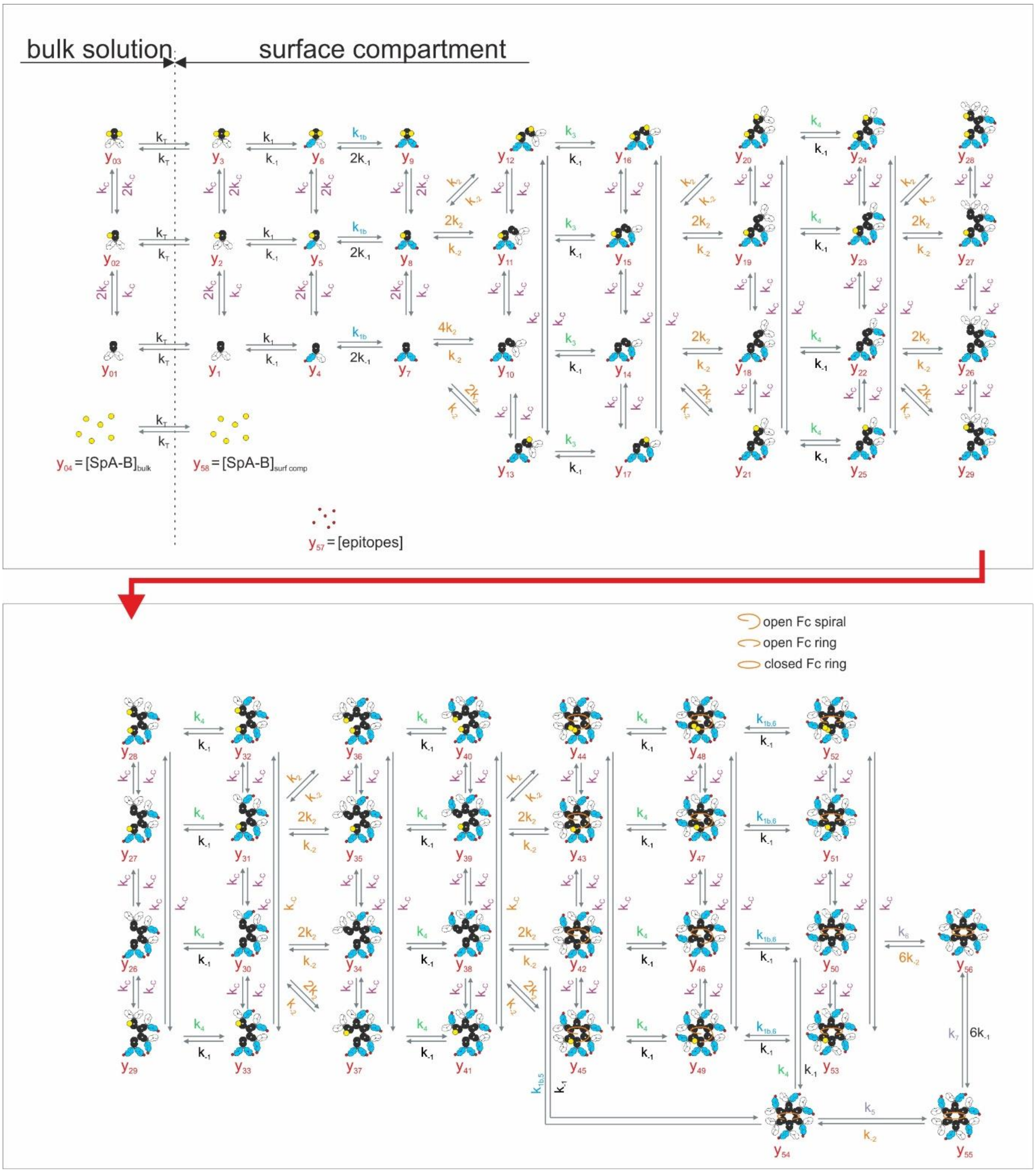


**Figure S1.** Kinetic model of antigen-dependent IgG oligomerization in the presence of an Fc binding and thus Fc-Fc interactions inhibiting protein like SpA-B. $k_C$ and $k_{-C}$ are the association and dissociation rate constants of the SpA-B – IgG Fc interaction.

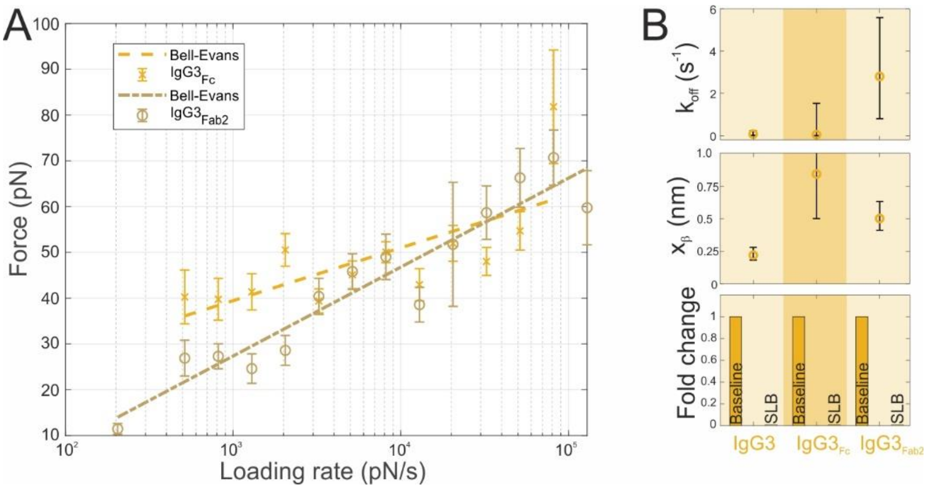


**Figure S2.** Single molecule force spectroscopy (SMFS) of IgG3 – IgG3 Fc and IgG3 – IgG3 Fab2 interactions. (A) Dependence of the most probable dissociation forces on the force loading rates (symbols) and least squares fits to the Bell-Evans model (solid lines). (B) Dissociation rate constant $k_{off}$ (upper panel) and distance from the bound to the transition state $x_b$ (middle panel) obtained from the fits in (A). Specificity controls employing bare SLBs.

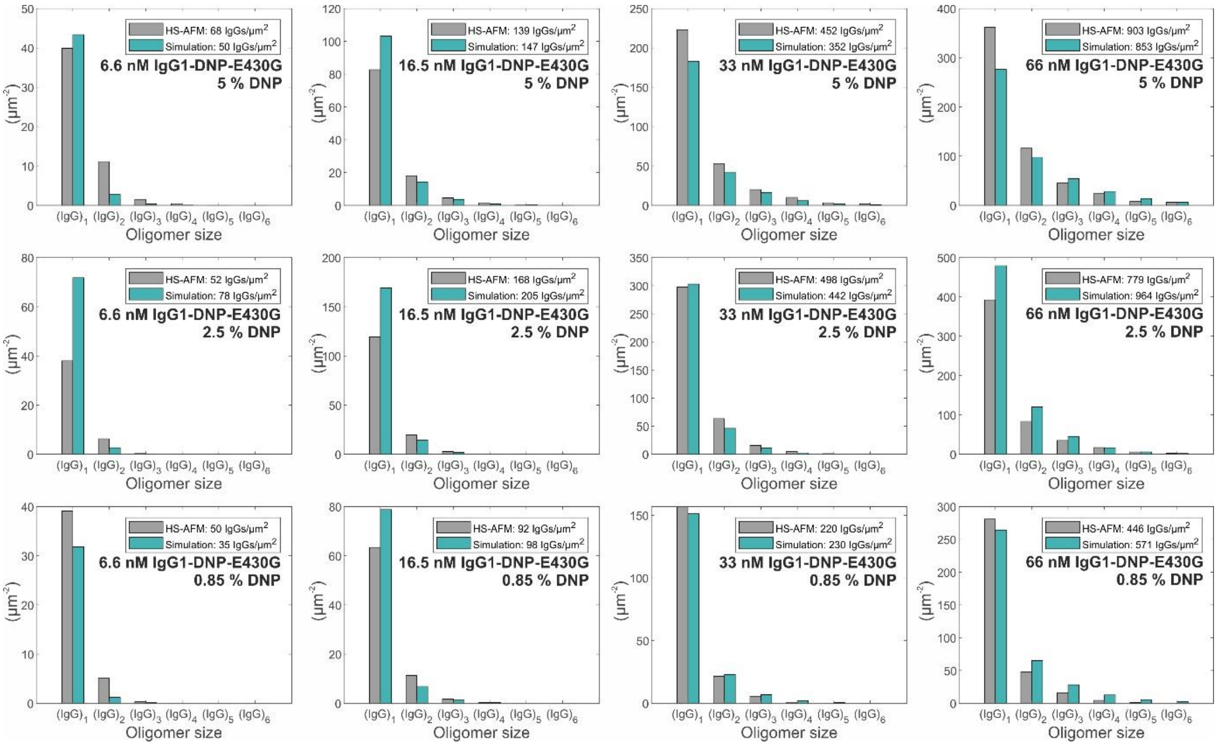

**Figure S3.** IgG1-DNP-E430G oligomer distributions obtained through 3 min incubation of 6.6 – 33 nM IgG1-DNP on SLBs containing 0.85 % DNP-DPPE (lower row), 2.5 % DNP-DPPE (middle row), and 5 % DNP-DPPE (upper row). Grey bars depict experimentally determined oligomer abundances; green bars represent a global fit (together with the data presented in Fig. 4, S4, and S5) to our kinetic model with parameters given in Table 1.

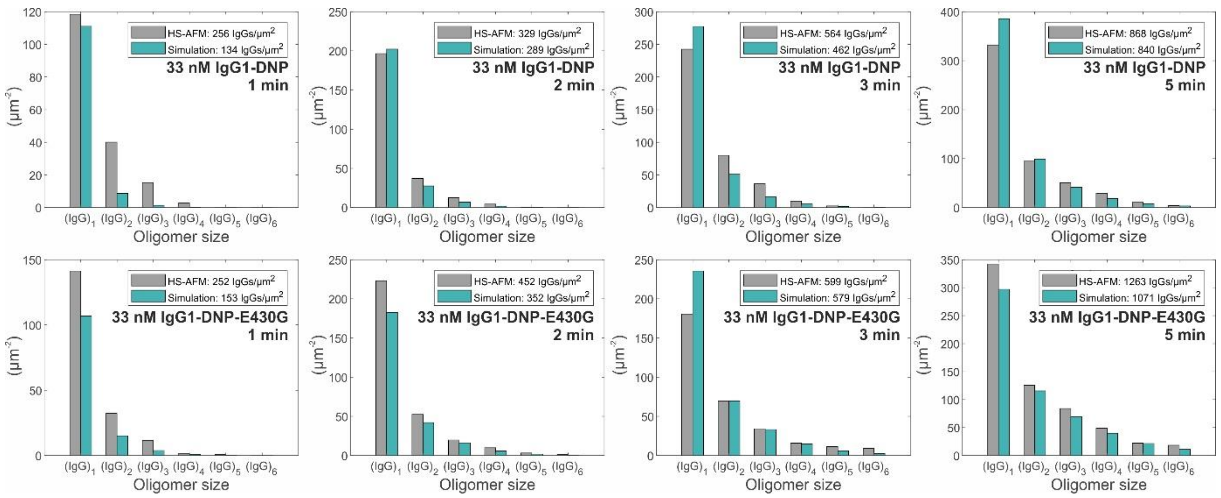


**Figure S4.** IgG1-DNP (upper row) and IgG1-DNP-E430G (lower row) oligomer distributions obtained through incubation of 33 nM IgG1 for 1 to 5 min on SLBs containing 5 % DNP-DPPE. Grey bars depict experimentally determined oligomer abundances; green bars represent a global fit (together with the data presented in Fig. 4, S3, and S5) to our kinetic model with parameters given in Table 1.

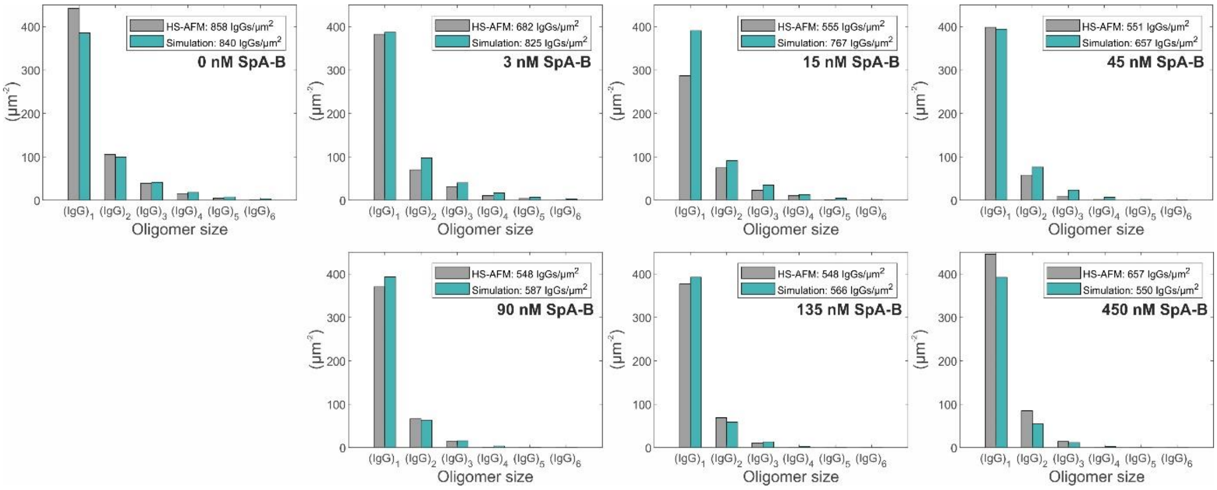


**Figure S5.** IgG1-DNP oligomer distributions obtained through incubation of 33 nM IgG1 in the presence of 0 to 450 nM SpA-B for 5 min on SLBs containing 5 % DNP-DPPE. Grey bars depict experimentally determined oligomer abundances; green bars represent a global fit (together with the data presented in Fig. 4, S3, and S4) to our kinetic model with parameters given in Table 1.

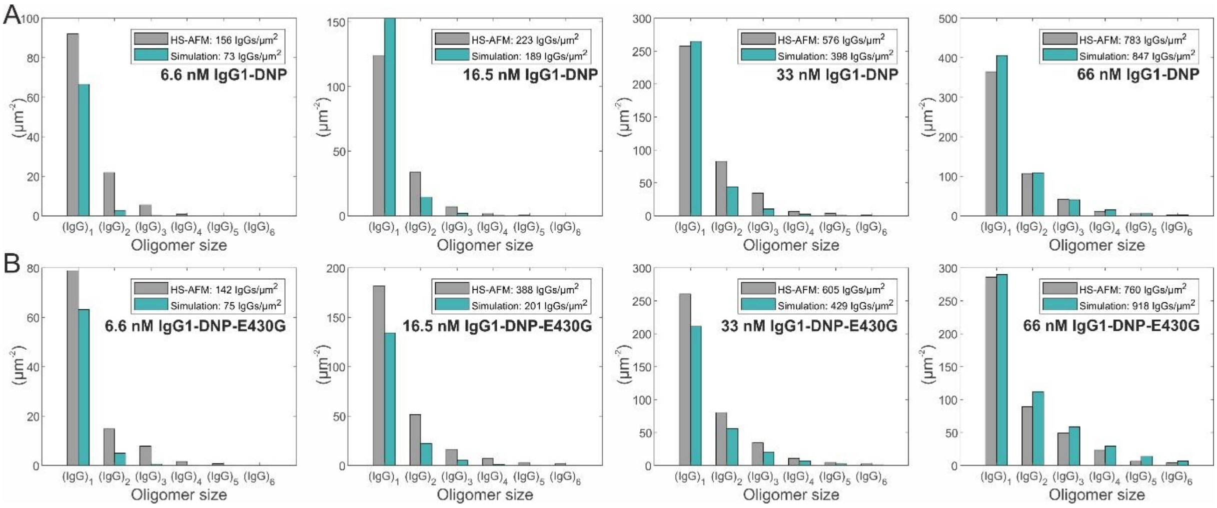


**Figure S6.** IgG1-DNP (upper row) and IgG1-DNP-E430G (lower row) oligomer distributions obtained through 5 min incubation of 6.6 – 33 nM IgG1-DNP on SLBs containing 5 % DNP-DPPE (upper row). Grey bars depict experimentally determined oligomer abundances; green bars represent a single parameter fit (mass transport rate from bulk into the surface compartment $k_T$) to our kinetic model with all other parameters fixed to the values given in Table 1.
